# Software for semi-automatic analysis of microscopic images of adhesion structures and protein colocalization in cells

**DOI:** 10.1101/2025.06.23.661040

**Authors:** Joanna Hajduk, Patrycja Twardawa, Zenon Rajfur

## Abstract

**Background and Objective:** Adhesion structures, such as focal and fibrillar adhesions, are protein complexes that mediate cell attachment to extracellular matrix via integrins. These structures participate in the mechanosensing process, transmitting forces from the cellular microenvironment to the cytoskeleton. The interaction between cells and their environment, along with the role of focal adhesion proteins, are areas of significant research interest. Accurate, quantitative analysis of adhesions structures in microscopic images is essential for advancing our understanding of the subject. However, the high variability and complexity of those structures in cells makes image analysis challenging, both in terms of subjectivity and time consumption.

**Methods:** We present a novel semi-automatic script for the detection of adhesion structures and analysis of their parameters, developed in MATLAB 2021a, to address the challenge of accurate image analysis of focal and fibrillar adhesions in cells.

**Results:** Our script offers a more time-efficient and less subjective alternative to manual analysis, while still allowing the user to retain control over the analytical process. It detects adhesion structures in confocal images and measures key parameters such as shape, orientation, and spatial distribution within cells, additionally providing visual label maps of the identified adhesion structures. The second add-on enables the calculation of correlation coefficients between two confocal microscopy image channels representing different stained cellular structures within the same cell, and generates visual colocalization maps, further enhancing the analysis of cellular architecture.

**Conclusions:** The presented open-source script offers a robust solution for the comprehensive, quantitative analysis of adhesion structures in microscopic images, based on user-defined analysis parameters. It is available online at https://github.com/patrycja-twardawa/FA-Colocalization.git.

## 1. Introduction

The cellular ability to sense and respond to various mechanical signals from the environment is fundamental for physiological function. This process, known as mechanotransduction, is facilitated by integrin-based adhesion structures, which connect cells to extracellular matrix (ECM) proteins. These protein complexes are involved in many crucial processes, including cell migration, proliferation and embryogenesis (1–3). Understanding the mechanisms behind mechanosensing and cellular responses to the mechanical properties of the microenvironment has been a focus of extensive research over last years.

Three types of adhesion structures (AS) can be distinguished, depending on their molecular composition, size, location within cell and maturation stage. Small, early-stage adhesion structures are called nascent adhesions. They are typically small (<1 μm), dot-like structure, formed at the edge of lamellipodium (1). Focal adhesions (FA) are more mature structures, larger than nascent adhesions and connected with actin stress fibers (1,4). They are composed of numerous proteins, such as talin, paxillin, vinculin, and signalling molecules, like focal adhesion kinase (FAK). Focal adhesions can further mature into fibrillar adhesions – elongated structures associated with fibronectin fibrils, which are typically found in the inner regions of the cell, closer to the nucleus. (5,6).

Traditional methods for quantitative analysis of adhesion structures, such as manual contouring and segmentation, have several drawbacks. They are time-consuming and therefore impractical for analyzing large datasets of cells containing numerous focal adhesions. This often leads to the analysis of only selected adhesions structures, a choice which may be subjective and reliant on personal judgment. On the other hand, automatic analysis approaches present challenges as well, due to diversity of adhesion structures in terms of size and shape, as well as varying quality of microscope images. Several software tools have been developed for at least partially automatic focal adhesion analysis. However, many of these tools have limitations: some are not readily accessible online, while others are tailored to specific imaging modalities or are difficult to use and modify. For example, Berginski and colleagues developed a system that analyses TIRF images of living cells, particularly useful for studies of adhesion structures dynamics (7,8). While freely available, the system allows modification of only a limited number of parameters and requires online processing on a server, making the analysis more time-consuming. Wurflinger et al. proposed an algorithm for FA segmentation and dynamic tracking (9). However, to our knowledge, this software is not open-source. Buskermole et al. designed an image analysis system for segmenting the actin cytoskeleton, nucleus and focal adhesions (10). While this program has various applications, it offers less parameter customization than our proposed solution.

In this paper, we propose a semi-automatic approach for the analysis of adhesion structures in microscopy images. Our method incorporates the efficiency of automatic processes such as ROI selection, image processing (contrast adjustment, denoising, background subtraction, filtration), segmentation, classification and labelling of structures, and quantification of their properties. Nevertheless, the user retains the ability to adjust segmentation methods, filtration criteria, and manually refine the resulting maps of structures by adding or removing objects and merging labels as needed. The parameters for automatic functions may be customized for various imaging conditions and cell or structure types. Additionally, for double-stained images, the script offers advanced features such as correlation coefficient calculations between two selected channels and the generation of colocalization maps, providing comprehensive insights into the spatial relationships between stained cellular structures.

## 2. Methods

### 2.1. Cell model

The algorithms for adhesion structure detection and colocalization assessment presented in this study were optimized and tested using the mouse embryonal fibroblast NIH/3T3 cell line. To evaluate the applicability of script to different cell types, the program was tested on the U2OS (osteosarcoma) cell line as well (Fig. S1). NIH/3T3 cells were cultured in Dulbecco’s Modified Eagle Medium with low glucose (Biowest), supplemented with 10% fetal bovine serum (FBS; Gibco) and 1% of Penicillin/Streptomycin (PS; Biowest). Cells were incubated in 37°C and 5% CO_2_ for approximately 20 hours on either glass-bottom dishes or polyacrylamide substrates. Substrates were prepared as described before (11).

### 2.2. Cell immunostaining and image acquisition

Cells were fixed in ice-cold methanol for 3 minutes, followed by washing with PBS and blocking with 4% BSA in PBS for 1 hour. Cells were then incubated overnight with mouse anti-talin-1 antibody (1:100; Bio-rad). Subsequently, they were incubated for 1 hour with Alexa Fluor 488 or 647 goat anti-mouse IgG (H+L) antibody (1:100; Invitrogen) and washed several times before imaging. Images were captured using a Zeiss LSM 710 confocal scanning microscope equipped with 40x oil objective (Plan-Apochromat 40×/1.4 NA; Zeiss).

## 3. Script description: tool for adhesion structures detection and colocalization analysis

The script was developed using MATLAB 2021a and is available as open-source software (URL: https://github.com/patrycja-twardawa/FA-Colocalization.git). Its functionality includes automatic adjustment of image properties, as well as segmentation and labelling of detected structures that display features of cellular adhesion structures, including fibrillar, focal, and nascent adhesions. The script also supports the generation of colocalization maps, which characterize the spatial relationships between all stained cellular structures (not only limited to those classified as adhesions) in two selected image channels. User retains full control over cell area and object selection and can influence feature extraction and classification by adjusting the available parameters. Alternatively, the entire process can be fully automated if the default parameters are deemed suitable. Simplified block diagram illustrating script function is showed in Figure 1.

**Fig. 1.**
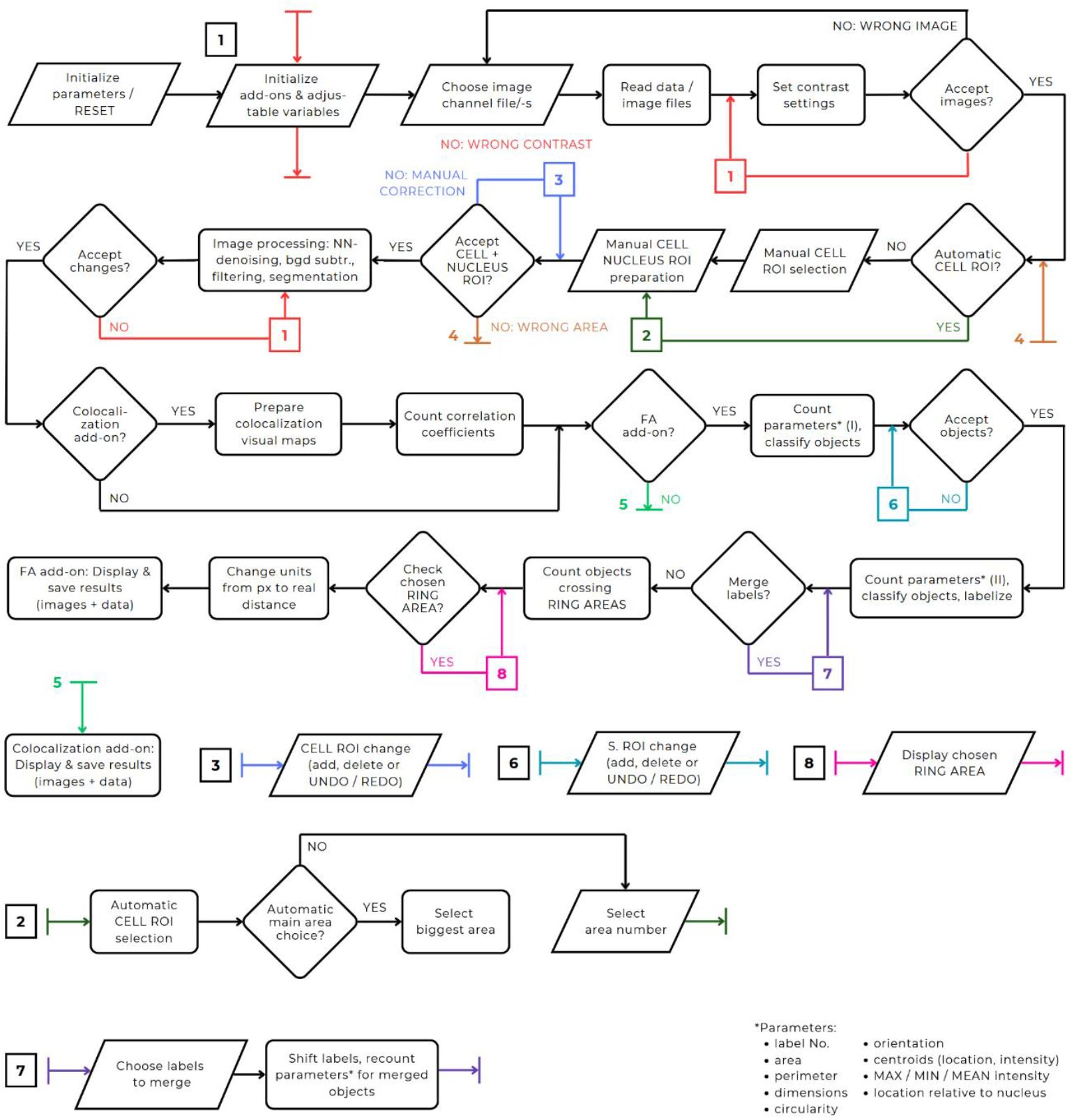
Simplified block diagram illustrating the functionality of the script, including the *Focal Adhesions* (*FA*) and *Colocalization* add-ons. The script is divided into independently executable sections, represented by rectangular blocks (automatic actions) and parallelogram blocks (mandatory user input). Diamond-shaped decision blocks indicate the recommended order in which the user should run the sections.

The script can be used directly on original images acquired via confocal microscopy, enhancing both their visual appearance and image properties. Although the script is suitable for analyzing any cell type, the default parameters are optimized for fibroblasts. When analyzing other cell types, additional parameter adjustments may be possibly required, based on testing and evaluation performed by the user.

A brief description of the sections is provided below. For more detailed information on script usage in specific cases, please refer to the Script Manual and Supplementary Materials.

### 3.1 Initialization of parameters and adjustable variables

In the first section of the script, essential parameters are initialized for both add-ons (*Focal Adhesions* and *Colocalization*). For most subsequent images, empty structures of appropriate size are preallocated to reserve memory in advance. Parameters marked as “*non-adjustable*” should not be modified or removed from the workspace by the user. All adjustable parameters are listed in Supplementary Materials (Tab. S1).

### 3.2 Choice of image files and initial preparation of image data

The next section of the script enables image file loading and contrast adjustment, allowing the user to view the image in a way that best supports tracking the operation of script. The selected file should contain a single channel of a confocal image if only the *Focal Adhesions* add-on is used, or two separately provided image channels if the *Colocalization* add-on, or both add-ons, are used. The image quality should be sufficient to clearly distinguish cellular structures (minor noise artifacts are acceptable). An image size of at least 2048 × 2048 px is recommended.

During this stage, additional file information is also read from the metadata of the provided image files, including the conversion rate from pixels to real-world units. If no pixel size information is available in the metadata, the user will be prompted to provide the conversion value manually.

Original confocal images may have low contrast settings, which can hinder the user’s ability to manually apply changes and fully benefit from the semi-automatic workflow. While the script operates by default with automated contrast settings, separate display settings are applied for better visualisation. If desired, the contrast used for display can also be applied to analysis via the “*adjustable parameters*” section. More robust contrast enhancements (e.g., lowering the upper intensity limit to increase the number of pixels treated as maximum intensity) may assist in detecting low-emission objects during segmentation. However, this may also increase the likelihood of including artifacts.

### 3.3 *Cell ROI* selection

The selection of the region of interest (ROI) refers to outlining a specific area in the image, which is represented by a binary map that stores information about the selection. The *cell ROI* refers to the area outlining the cell body and its interior. It is extracted after the image and metadata are read from the provided original file, and the method (automatic / manual) is selected based on user’s choice declared via adjustable flag value. Automated *cell ROI* selection is based on the analysis of local intensity densities. A moving mask centred on each pixel scans the image, and if fewer than 65% of the pixels within the mask exceed the default brightness threshold, the central pixel is excluded from the selected area. Conversely, if the threshold is met or exceeded, the central pixel is included. Once the entire image has been processed, the resulting binary map is refined using edge smoothing and hole filling.

The result of automated cell ROI selection may include several detected objects. If this occurs and if the user has enabled object selection for the *cell ROI*, a labelled map of the detected objects is displayed. The user can then select the desired region by entering the corresponding object label number in the console window. This feature allows individual analysis of multiple cells captured in a single image. If object selection option is disabled, the biggest object will be selected automatically.

Automatic *cell ROI* may be manually adjusted or completely replaced by the user. *Cell nucleus ROI* selection is optional and may only be done manually.

### 3.4 Image processing and segmentation, *Segmentation ROI* selection

After the *cell ROI* has been selected, the image undergoes further processing to enable the extraction of cellular structures. The processing workflow consists of several steps, which differ between the *Colocalization* and *Focal Adhesions* add-ons. The *Focal Adhesions* add-on includes an additional step, based on Hessian-based Frangi Vesselness filtering method to enhance the intensity of fibrillar structures (12,13). This step is omitted in the *Colocalization* add-on to allow for the inclusion of a broader cell area, beyond only adhesion structures. The earlier processing steps are shared by both add-ons and are executed in succession: image denoising, background subtraction, and filtering.

Denoising is performed using a pre-trained *DnCNN* network (12). Background subtraction is carried out based on morphological operations (mathematical morphology) (12,14). Various filter types can be selected, such as bilateral, Gaussian and Savitzky-Golay filters, with selectable smoothing levels (12,15,16). By default, a bilateral filter is applied.

After image processing, segmentation is applied within the previously extracted *cell ROI* (Fig. 2B). The same ROI is used for both script add-ons. Users can choose between adaptive and global segmentation methods. The *Colocalization* add-on performs segmentation on the sum of two normalized image channels provided by the user, using its separate gain values. Global segmentation with automatic thresholding via Otsu’s method is recommended for fibroblasts (12). However, the user may also manually specify a threshold or select among other provided local segmentation methods if desired. In cases where structures are faintly visible in the original image, local segmentation may yield better results, though it may also introduce artifacts, particularly near the cell nucleus.

**Fig. 2.**
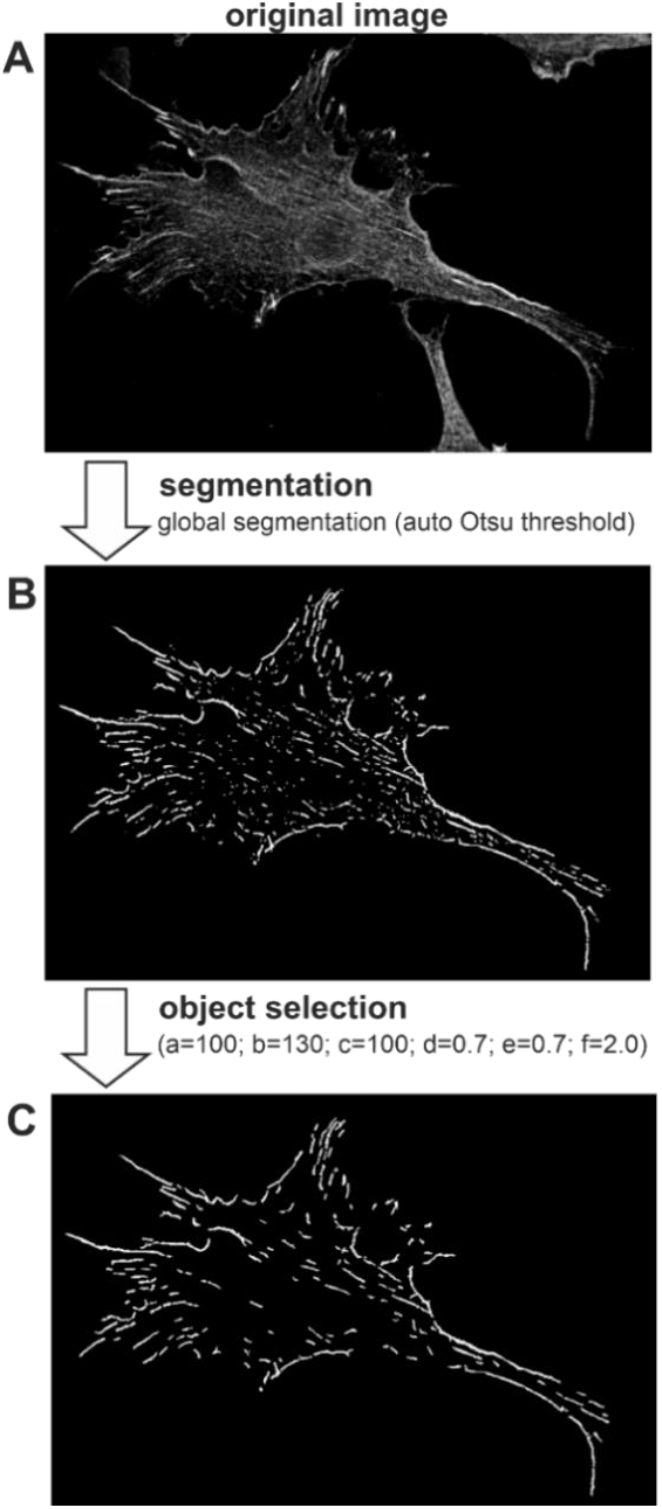
An exemplary result from the segmentation (*B*) and classification (*C*) stages of image analysis, performed on a cell image acquired via confocal microscopy, following the initial processing stage executed by the script (*A*). Classification parameters: *a* – minimum number of pixels assigned to fibrillar objects with intensity above the set threshold; *b* – minimum number of pixels assigned to larger circular objects with intensity above the set threshold; *c* – minimum number of pixels assigned to objects, regardless of other factors; *d* – circularity threshold (above – objects considered more circular, below – object considered more fibrillar); *e* – intensity threshold; *f* – minimal *major-to-minor dimension ratio* for more fibrillar objects, to be considered accepted as adhesion structures.

The output of segmentation is a binary map of objects, separately for two add-ons and for cell nucleus areas. For the *Colocalization* add-on, this map represents all cellular structures visible in both channels. For the *Focal Adhesions* add-on, it includes detected adhesion structures (the target objects) along with potential artifacts. It is recommended not to manually exclude unwanted objects prior to classification; instead, all potential cellular adhesion candidates should be retained. Undesired objects should be filtered out later during classification stage. Users can manually modify the *segmentation ROI* during later stages (see section 3.6.3) using the *Focal Adhesions* add-on, whereas for the *Colocalization* add-on, modifications to the ROI encompassing cellular structures are available only through the “*adjustable parameters*” section of the script.

### 3.5 *Colocalization* add-on operations

Colocalization add-on analyses the relationship between two selected image channels representing different stained cellular structures/proteins in the same cell by the analysis of whether stained structures in both channels share matching spatial positions. The script enables an automated image processing procedure for the purpose of preparing colocalization maps and evaluating correlation parameters.

#### 3.5.1 Colocalization maps preparation

Two analysed proteins may not exhibit the same emission intensity, depending on used antibody or on measurement parameters (gain, laser power etc.), resulting in potentially lower pixel values in one of the channels. Thus, it is essential to first normalize each channel to its respective maximum value. This is a necessary initial step performed during earlier stages of image processing. It should be noted that, after normalisation, image intensities are relative to their maximum values and enhanced, which may lead to the appearance of noise artifacts in channels with very low maximum fluorescence intensity. All colocalization maps operate within the area defined as the cellular structures area, determined by the segmentation step specific to the *Colocalization* add-on.

To assess the relationship between intensities in the two selected channels, four colocalization maps are generated, each representing different properties of the channel interactions. These maps are: *Sum Map, RGB Overlay Map, Subtraction Map*, and *Hue Map* (Fig. 3, Fig. S2).

**Fig. 3.**
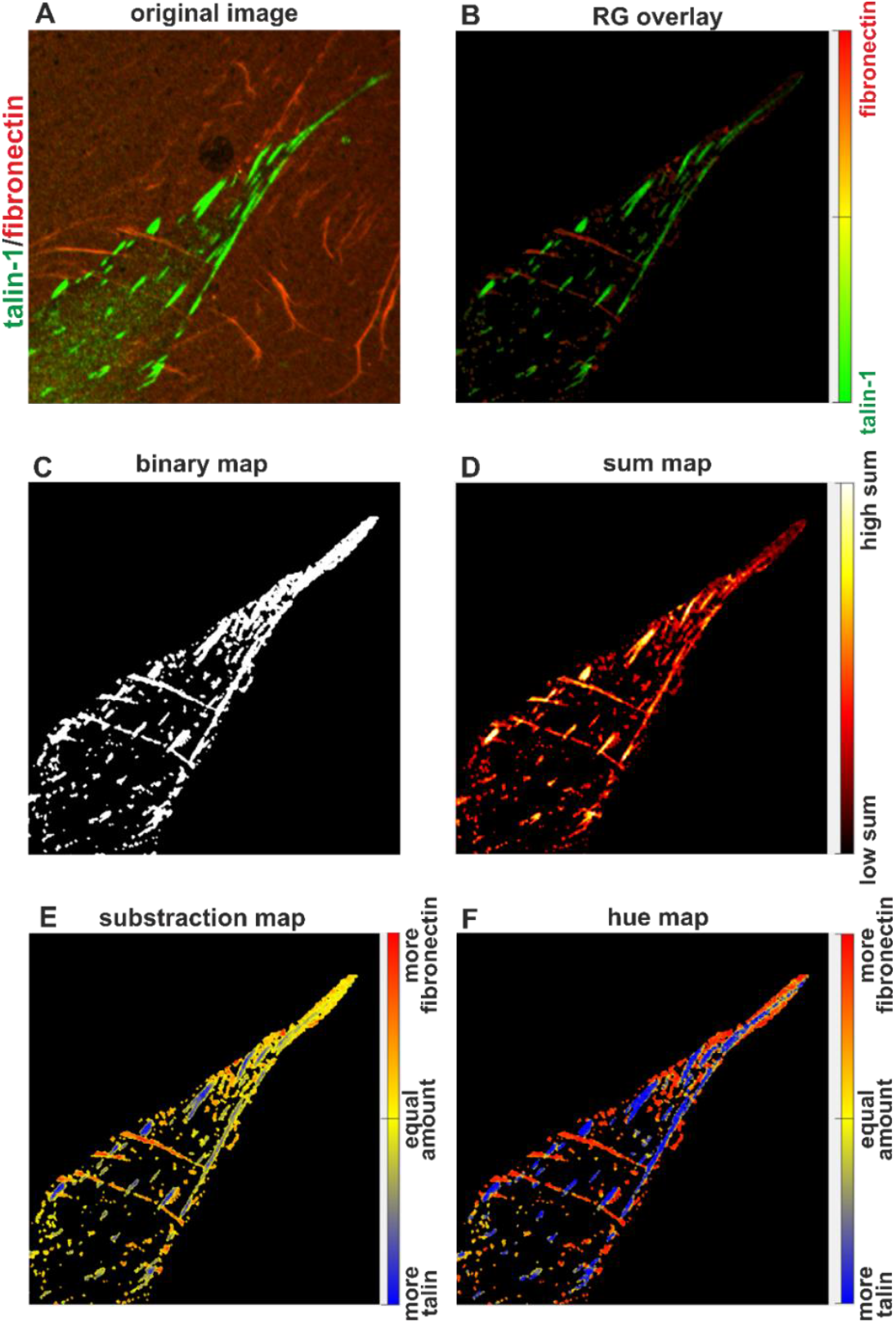
Exemplary results of colocalization maps preparation. (*A*) Original image with adjusted contrast (green – Talin-1; red – Fibronectin), shown as an overlay of both image channels, including areas outside the identified cellular structures. (*B*) *RGB Overlay Map*. (*C*) Binary map representing structure areas found in both provided image channels, after segmentation. (*D*) *Sum Map*. (*E*) *Subtraction Map*, with labels adjusted for the specific case (label names can be modified in the “*adjustable parameters*” section of the script). (*F*) *Hue Map*, with label names assigned for the specific case.

*Sum Map* displays the normalised sum of intensities from both channels at locations corresponding to stained cellular structures. It does not differentiate between channels but focuses on their combined intensity, which reflects the density of stained structures at a given location. *RGB Overlay Map* is a simple overlay of both channels using one of the three colors channels (red, green, or blue), as selected by the user – regardless of the original fluorescence color. It provides a visual merge of the two channels in a single image. *Subtraction Map* visualizes the difference between the normalised intensities of the two image channels. As the result, values greater than zero (indicating higher intensity in the first channel) are scaled and mapped to the range (0.5, 1]; values less than zero (indicating higher intensity in the second channel) are scaled to the range [0, 0.5). Equal intensities in both channels are assigned a value of 0.5. This map highlights which channel dominates at each pixel relative to their combined intensity and also emphasizes low-intensity differences in dim regions – representing such areas with vivid colours, unlike the *RGB Overlay* or *Sum Map*, which may render them as indistinct background. *Hue Map* displays the relative contribution of each channel at each pixel location, regardless of absolute intensity. For instance, if both channels have low intensity at a given pixel but one channel is still more prominent, the *Hue Map* will display a colour indicating that dominance – even though the absolute intensity is low. This approach differs from the *Sum Map, RGB Overlay*, and *Subtraction Map*, which are all influenced by total signal strength.

#### 3.5.2 Correlation coefficients calculation

In addition to the visual representation of colocalization maps, numerical results representing the correlation between channels are provided in the form of correlation coefficients, calculated for two-dimensional maps at locations corresponding to the identified cellular structures. Two types of coefficients are computed: Spearman’s rho and Pearson’s linear correlation coefficient – each calculated separately for the entire map and for the nucleus region (specifically within the *segmentation ROI* overlapping with the *cell nucleus ROI*) (12).

To assess these correlation coefficients, the two-dimensional image matrices are flattened into one-dimensional data vectors for each image channel. A strictly linear relationship between the channels is not expected; rather, the desired outcome is the detection of monotonic co-fluctuations between the two vectors – assuming the stained structures are spatially colocalized. Given the anticipated nonlinear relationship between channels, Spearman’s rho is the recommended measure for assessing correlation.

It is important to note that, for any of the provided methods to be valid, the image chromatic aberration should be corrected. The script does not perform realignment of the provided image channels, so the user must ensure proper alignment beforehand.

#### 3.5.3 Data display and saving for *Colocalization* add-on

Colocalization maps are displayed immediately after generation, while the corresponding correlation coefficient data are stored in a dataset ready for export. Running the final section results in saving the images in both FIG and TIFF formats, and the correlation coefficient data in an XLSX spreadsheet, all within the folder containing the first provided image channel.

### 3.6 *Focal Adhesions* add-on operations

The main add-on of the script is designed to identify objects that may correspond to cellular adhesion structures, compute their properties, and provide visual representations showing the locations of the detected structures. The quality of object segmentation and classification depends heavily on the processing stage, which can be influenced by the user depending on the specific case and related requirements (see section 3.4). Adjustable parameters and options are listed in the Supplementary Materials (Table S1).

#### 3.6.1 Object parameters and descriptors estimation

The script calculates the basic parameters of the detected objects, considered as adhesion structure candidates. These parameters include object dimensions (*area, major* and *minor dimensions, major-to-minor dimension ratio, perimeter*), *centroid location* (with vertical axis *Y* and horizontal axis *X* coordinates, in that order, in consecutive columns), *circularity, orientation*, and intensity-related data (*minimal intensity, maximal intensity*, and *mean intensity* values, *weighted centroid*). Additionally, the resulting table records whether objects are located above the cell nucleus area, based on the *cell nucleus ROI* provided by the user. Examples illustrating data presentation in the results table are provided in the Supplementary Materials and the script manual, while a short description has been provided below.

The area (*A*) refers to the total area covered by an object, based on the number of assigned pixels. The *major* and *minor dimensions* refer, respectively, to the longest and shortest diameters of the object. It should be noted that if an object consists of a spiral structure, these diameters may pass through background areas separating different parts of the object, such as when an outer fragment envelops a central region. The *major-to-minor dimension ratio* refers to the result of dividing the *major dimension* of object by its *minor dimension* value. The perimeter (*P*) of object is estimated by calculating the distance between each adjoining pair of pixels, using the 8-neighborhood connectivity. The *centroid location* is expressed for each object as a two-element vector with mean rounded coordinate values, provided in separate columns (first the vertical *Y* axis representing image matrix rows, then the horizontal *X* axis representing image matrix columns). *Circularity* (*form factor, FF*), ranging from 0 to 1, is calculated using the following formula:

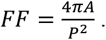

For a perfect circle, in theory, *FF* = 1. In practice, circular objects can reach *FF* values greater than 1 due to digitization errors, particularly in the case of small objects. Since the analyzed images are inherently pixel-based, this error cannot be completely eliminated in the calculation. However, the *FF* parameter is widely used, and the impact of digitization error becomes negligible for larger objects composed of many pixels. Given the significant differences in *FF* values typically observed between fibrillar and circular structures, the parameter remains effective for filtering out predominantly circular objects.

Orientation (*O*) is calculated as the angle between the horizontal *X* axis and the major axis of an ellipse that shares the same second moments as the object. The value of *O* ranges from -90° to 90°. There are also measures related to object emission intensity, where intensity for each pixel is normalised to the maximum intensity value within the *cell ROI*. This normalisation stretches the scale, enabling a consistent map display across different cell images. The primary intensity parameters include *minimal* (*I*_*min*_), *maximal* (*I*_*max*_) and *mean* (*I*_*mean*_) intensities of objects. Additionally, the intensity-weighted centroid (*I*_*wc*_) is calculated, representing the central point of the region based on both pixel location and intensity value.

#### 3.6.2 Classification of objects

The binary map after segmentation may include both adhesion structures and artifacts. Therefore, classification is a necessary step to differentiate between these two types of structures (Fig. 2C). In the first step, parameters are calculated for all segmented objects. Based on these parameters, the classifier then excludes objects that do not meet the criteria for adhesion structures. These criteria include: object circularity, area, emission intensity, and the ratio of dimensions. Objects accepted as adhesion structures should exhibit characteristics associated with specific types of adhesions:

- Fibrillar adhesions: long (>5 μm) fibrillar objects with medium to high staining intensity;
- Focal adhesions (FA): medium-sized (1–5 μm) fibrillar or ellipsoidal structures with medium to high staining intensity;
- Nascent adhesions (NA): small (0.5–1 μm), dot-like structures with medium to high staining intensity (10).

Excluded structures typically include small circular and non-circular objects with low emission intensity, very small non-fibrillar objects, and the smallest objects with very low emission intensity. These often result from artifacts introduced during filtering and smoothing, where typical noise from confocal imaging aggregates into larger structures.

Users can modify the area and intensity thresholds for circular and fibrillar objects. It is recommended to set the threshold for circular object area higher than threshold for fibrillar objects to avoid excluding potential candidates for nascent adhesions. While smaller size and circular shape are expected features of nascent adhesions, elongated objects of similar size are more likely segmentation artifacts. Focal adhesions and fibrillar adhesions typically appear as larger structures, reflecting their role and developmental stage.

If objects are manually added to the binary map after segmentation by the user, the classifier offers an option to prevent their exclusion via the flag available in “*adjustable parameters*” section, even if they do not meet the classification criteria. This feature supports the inclusion of small objects the user considers likely nascent adhesions. By default, this option is enabled, but users can disable it to apply the same classification criteria to both detected and manually added objects. If any objects added manually by the user were accepted by overriding the classifier rules, an additional column will appear in the dataset to mark those structures.

#### 3.6.3 *Segmentation ROI* adjustment

User can apply modifications to the resulting *segmentation ROI* after classification, which represents the areas attributed to objects selected as adhesion structures from among all candidates. At this stage, individual objects are not yet recognized by assigned labels. Modifications to the *segmentation ROI* are performed similarly to manual corrections made to the *cell ROI*. Available options include adding and deleting objects, with the possibility of undoing and redoing actions via the “*UNDO / REDO*” subsection of the script, which stores the memory of one user action backward or forward.

#### 3.6.4 Final object parameter recalculation and classification, labelling

Final recalculation is performed after user accepts the *segmentation ROI* and after any eventual manual modifications to its structure have been made. The parameters of the objects are re-estimated using the final version of the *segmentation ROI*, after which labels are assigned to the objects, with numbers increasing from the left side of the image to the right. Each object receives a unique number, which is used to represent it in the numeric data table and visual label map (Fig. 6C). By comparing the visual label map with the numeric data table, user can identify the parameters corresponding to specific objects.

#### 3.6.5 Label merging

Software users can merge several separate objects into a single larger object after the classification stage (Fig. 4). Occasionally, stained fibrillar structures are captured with lower emission intensity in certain fragments, resulting in their segmentation as separate objects due to breaks between them. Using the label map provided after classification, users can manually merge such structures if they determine that the fragments belong to the same object.

**Fig. 4.**
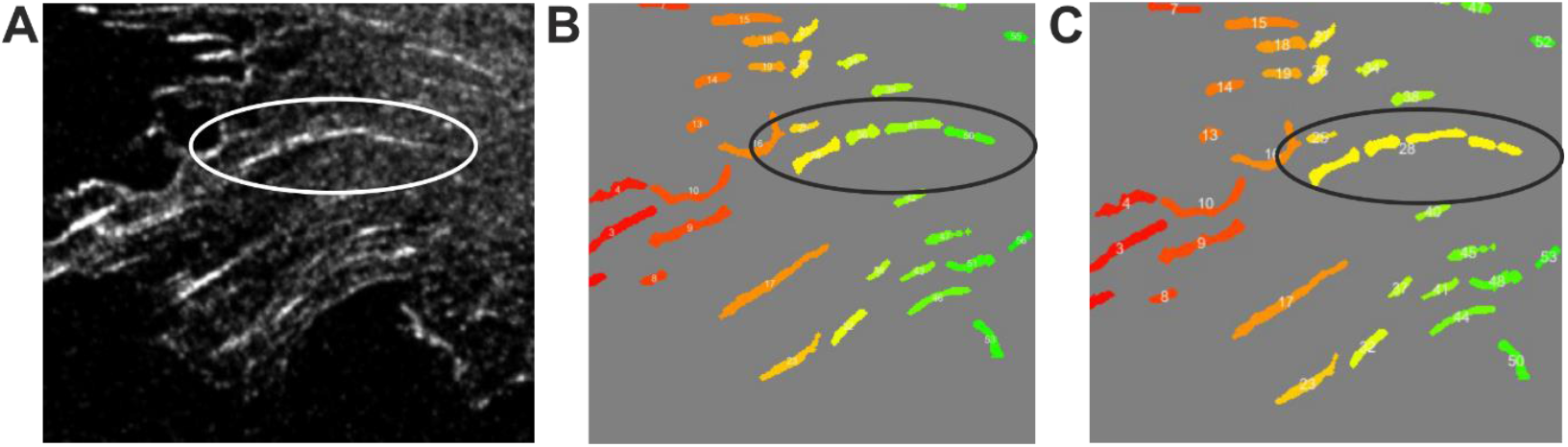
Exemplary illustration of the label merging stage. In the original image (*A*), it is apparent that the four objects identified after segmentation (*B*) are parts of a single elongated structure. Therefore, they were manually merged and assigned a single label (*C*). This stage was performed manually, and the decision was made by the user.

To merge objects, users can outline the desired objects or parts of them (including only a portion of a specific object is sufficient for it to be fully included). All objects within the outlined area will be relabelled as a single structure (Fig. 4). The new object label will correspond to the lowest label among the merged objects, while the labels of all subsequent objects will be shifted and reassigned to maintain sequential labelling. Once merging is complete, the parameters for the new object are recalculated based on the scores of the original objects, as described in the script manual. While labels can be merged, they cannot be split at this stage, because merging does not alter the binary map obtained after classification, whereas splitting would.

Merging typically involves connecting parts of an object separated by areas of very low intensity. Attempting to manually fill these gaps could lead to arbitrary decisions rather than the clear identification of structures. The merging feature is designed to minimize such arbitrariness by allowing users to connect only the visible parts of objects into a single structure. If users prefer a different approach, they can manually draw objects during the earlier stages, prior to classification, which serves as an alternative to merging.

#### 3.6.6 Division of cell area into *ring areas* and related calculations (change placing in code)

The cell area, based on the *cell ROI*, can be divided into *ring areas* (Fig. 5A) in order to track either the gradual change in intensity or the density of object placement relative to specific cell regions. The first “*ring*” is defined as the region closest to the cell edge, with a default width of 1 μm (a user-adjustable parameter). The ring width can be adjusted by the user but cannot exceed the minor dimension of the object. Subsequent rings are extracted consecutively toward the cell interior until reaching its midpoint.

**Fig. 5.**
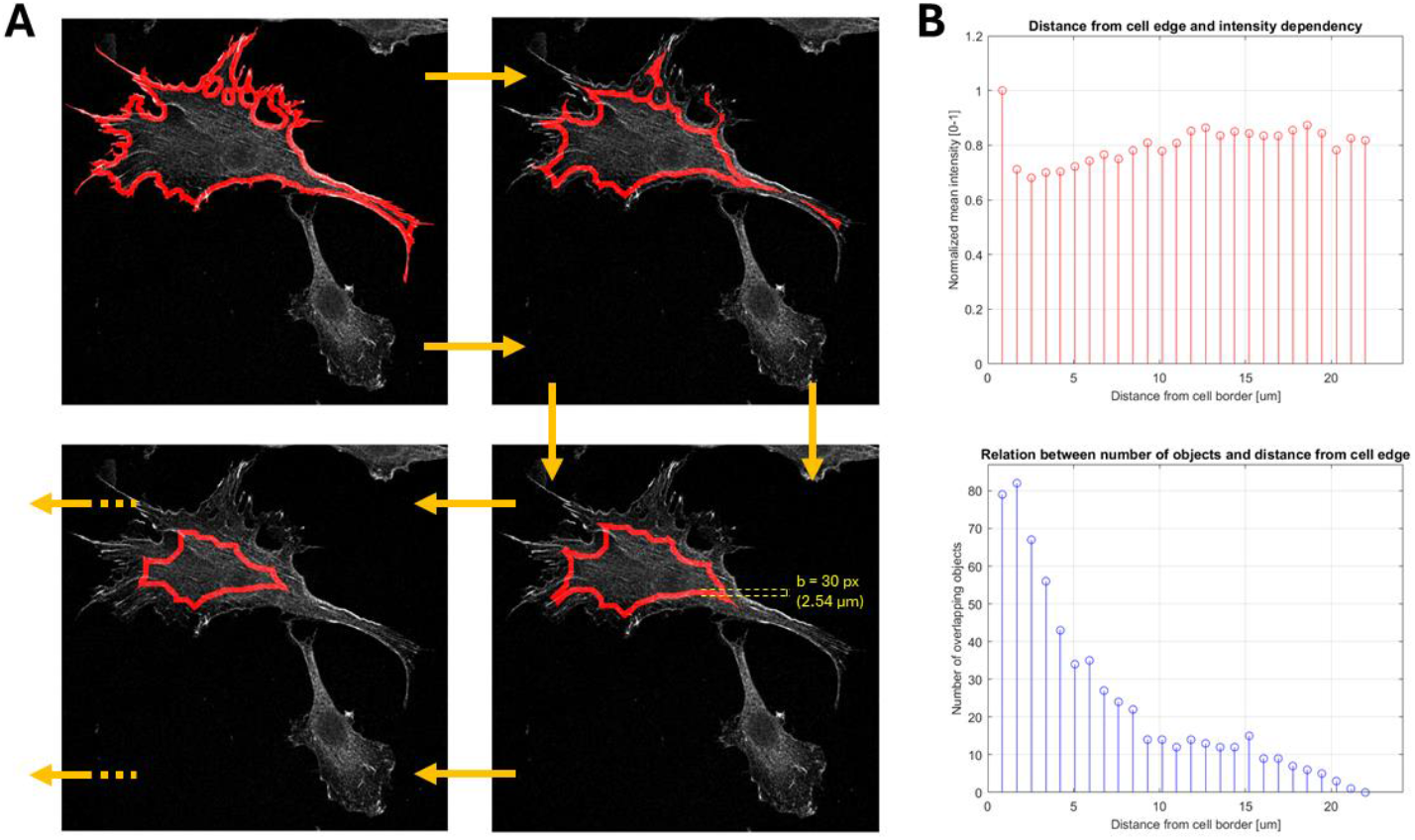
Examples of extracted *ring areas*. (*A*) Image showing the outer *ring area* overlaid on the cell image, with the *ring area* width defined by the user (here: 30 px). (*B*) Charts illustrating: (top) *mean intensity* level in consecutive *ring areas* (moving from the cell’s outer edge toward its interior) as a function of distance from the cell edge; and (bottom) the *number of overlapping objects* (based on the label map) as a function of distance from the cell edge.

**Fig. 6.**
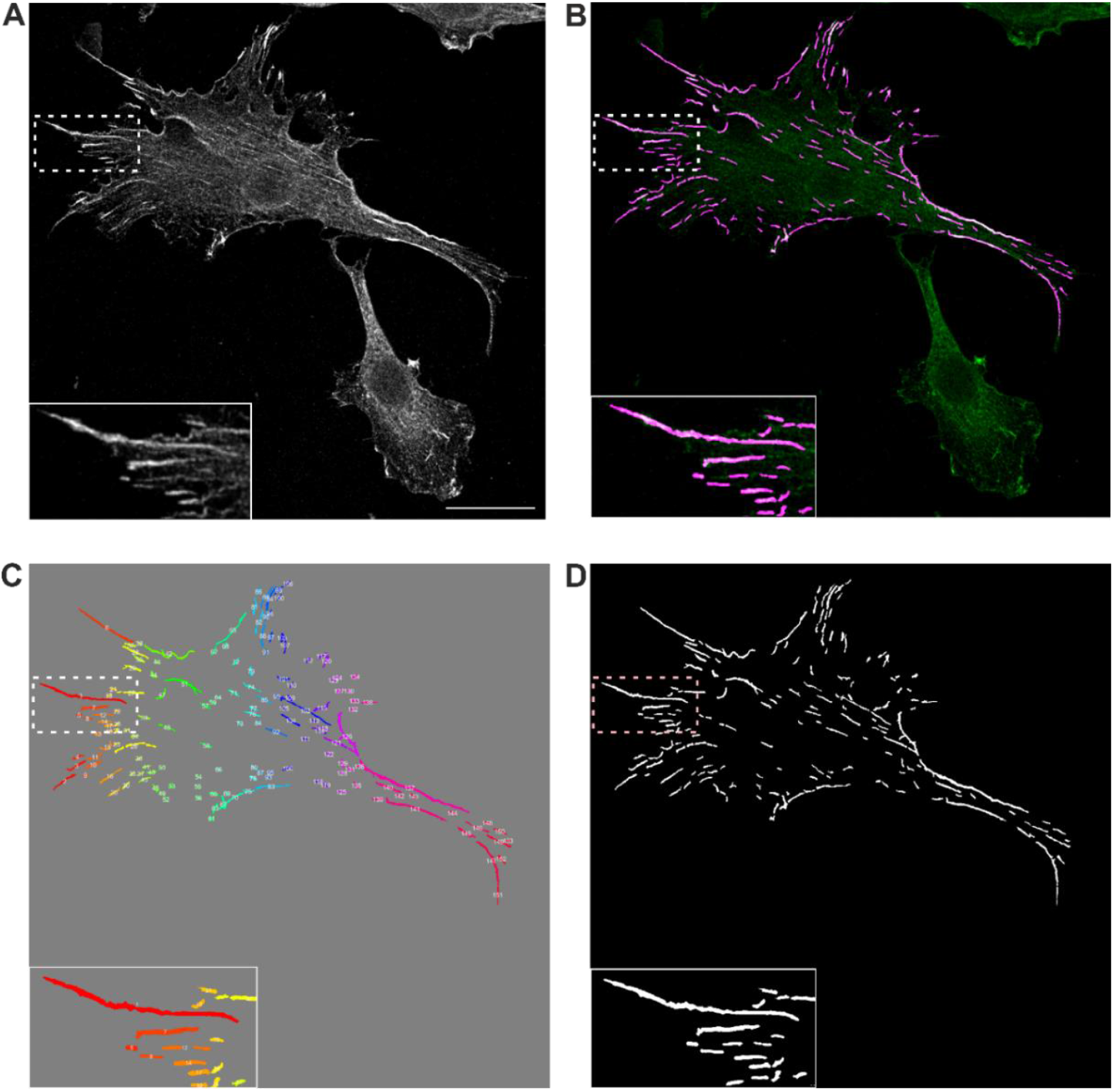
Exemplary result of segmentation and labelling of adhesion structures in an NIH/3T3 cell seeded on glass. (*A*) Original image of Talin-1 in an NIH/3T3 cell with adjusted contrast. Scale bar: 30 μm. (*B*) Overlay of the detected and classified adhesion structures (*segmentation ROI* marked in purple) on the original image channel (green). (*C*) Identified adhesion structures with assigned labels. Label colours are assigned randomly. (*D*) Binary map of the detected adhesion structures, without label division, after classification – representing the area of the final *segmentation ROI*.

The *ring areas* are generated by eroding the object map with a circular structural element (4-neighborhood). Parameters set counted for each *ring area* consists of the *mean intensity, mean normalised intensity, minimal intensity, maximal intensity, total area, distance from cell border*, and *number of objects overlapping the ring area* (count after 2^nd^ estimation, move!).

All parameters are calculated for the entire *ring area*, which includes not only the regions covered by segmented objects but also the spaces between them, located within the cell body (*cell ROI*) and within the specific *ring area*. The *mean, minimum*, and *maximum intensity* values correspond to statistical intensity properties computed across all pixels within the given *ring area*. The *total area* refers to the number of pixels attributed to the *ring area*, which is later converted to real-world units (see section 3.6.7). The *distance from cell border* refers to the distance between the outer edge of the *ring area* and the cell contour. The *number of objects overlapping the ring area* indicates the number of identified adhesion structures that are fully contained within, intersect with, or partially overlap the analysed *ring area*.

The software is capable of plotting the mean intensity as a function of distance from the cell edge, with units corresponding to increasing multiples of the ring width, thus allowing assessment of intensity changes toward the cell interior (Fig. 5A). Another type of analysis counts the number of labelled objects overlapping each consecutive ring area, providing information on the density of object placement relative to the cell area they occupy (Fig. 5B). Combined data from both charts can offer insights into cytoskeletal build density and structural organization across different cell regions (e.g., filopodial, lamellipodial, and internal areas).

The software also supports hole detection within the cell area based on user preference. When this feature is enabled, ring areas will wrap around holes without crossing them during extraction. In this context, holes are defined as external background spaces fully enclosed by the cell body. If hole detection is disabled, any blank spaces completely surrounded by the cell body are treated as integral parts of the cell and included in intensity measurements; this choice will also result in a visible change in the displayed *cell ROI*. For the hole detection mode to function properly, the provided *cell ROI* must include these internal blank spaces, meaning that the spaces must be manually deleted from the *cell ROI*, as automatic extraction will otherwise fill the holes detected within the cell body.

#### 3.6.7 Recalculation to real units

After all subsequent calculations are completed, the script recalculates the values to real-world units instead of pixel units, using the pixel size obtained either from the file metadata or provided by the user during the file reading stage. The default real-world unit after conversion is microns (μm). If the section responsible for unit recalculation is not executed, the resulting values will remain in pixel units.

#### 3.6.8 Data display and saving for *Focal Adhesions* add-on

The numeric results are stored in the temporary data structure prepared for saving, while maps providing the visual information are displayed in separate windows. Resulting images consist of the processed original image (Fig. 5A), binary maps (map of fibrillar objects, objects excluded by the classifier), the overlay of the image channel and *segmentation ROI* (Fig. 5B), label maps (the object contour map with label numbers, maps of found objects with coloured labels, showing separately the cell, cytoplasm, and cell nucleus areas, Fig. 5C), and charts regarding ring areas data visualisation (chart of fluorescence intensity as a function of distance from the cell edge, chart displaying the count of adhesion structures as a function of distance from the cell edge, Fig. 4B).

All generated images and charts are automatically saved in both FIG and TIFF formats in the same directory as the original input image for *Focal Adhesions* add-on. The quantitative datasets obtained for adhesion structures alongside the parameters describing the *ring areas* are saved to a spreadsheet in XLSX format.

## 4. Discussion

The script enables automated detection of adhesion structures in confocal microscopy images of stained cells and provides tools for analyzing the spatial colocalization of detected structures. Two add-ons are available, each of which can run independently. Among the many possible approaches, our proposed solution combines the generation of both numerical data and visual maps, supports parameter customization, and allows for fully automated operation when appropriately configured for a specific use case – streamlining the analysis process and saving time. The software is fully open-source, ensuring broad accessibility.

Presented tool calculates multiple parameters for cellular adhesion structures, offering detailed and versatile output for various research applications. Although the script supports automated processing, it is highly adaptable, giving users precise control over the analysis workflow. Users can modify a wide range of parameters, including segmentation methods and preferred object sizes, to suit specific experimental conditions. Alternatively, default settings can be used, significantly reducing the time required for parameter tuning and image analysis – especially after initial manual adjustment, which can be applied to future analyses of similar images. Manual corrections – such as merging labels, adding or removing objects, and adjusting ROIs – can be performed at specific script stages where such modifications may be required, providing a balance between automation and user-driven refinement. This hybrid approach ensures both accuracy and flexibility.

In addition to adhesion structure analysis, the software supports the quantification (e.g., by calculating correlation coefficients) and visualization of colocalization between two stained proteins via the *Colocalization* add-on. This is achieved through multiple approaches that focus on different aspects of the relationships between channels, thereby enhancing the functionality of software and its potential applications.

It should be noted that users should be familiar with biological image interpretation and refer to the script manual, adhering to the recommended implementation guidelines to ensure valid and reproducible results. The reliability of the results and the appropriate use of script options depend heavily on the user’s knowledge and interpretation.

There are several limitations in the current implementation that should be considered before performing the final statistical analysis. Many potential issues, also among those mentioned below, can be mitigated by using high-quality images; however, since this may not always be possible, the script is designed to operate on images containing noise and small object artifacts – provided they are handled appropriately and with user awareness. Users should also note that in poorly stained or darker image channels, the normalization performed by the script may result in disproportionately high mean intensity values compared to better-stained samples or other well-stained image channels.

One of the most critical factors affecting the accuracy of results when using the *Colocalization* add-on is the requirement for precise spatial alignment of the provided image channels. If this condition is not met, all the calculated correlation coefficient values will be incorrect, and any statistical analysis based on such results cannot be considered valid.

Users should take into consideration also a few other issues, when working with the script. Automatic ROI detection is not reliable for closely positioned or touching cells, as both may be included within a single outline. However, the ROI can still be extracted automatically, and any excess cell area can be manually removed by the user. Fiber merging must be performed manually, as the current version of the software does not support automatic merging of segmented structures. Users should also be aware of how object parameters are recalculated after label merging. The area (*A*) becomes the simple sum of pixels from the merged objects. The orientation (*O*) may change due to updated second-moment calculations for the newly combined structure. The perimeter (*P*) typically increases, as the software connects separate objects using a thin line (visually zero-width but still contributing to the *P* value), which significantly affects the circularity (*FF*). This connection-based approach was considered fair, as it links structures without adding artificial components beyond the visibly merged objects. As a result, data for merged objects is inherently prone to increased errors in *P* and *FF*, due to the presence of imaging discontinuities that the script cannot correct within a fair modelling framework.

The program may also encounter difficulties with large background artifacts, particularly near the nucleus, where distinguishing true structures from noise is more challenging. To address this, the script includes options for manual ROI adjustment, enabling users to remove such artifacts when necessary.

## 5. Conclusions

The proposed script provides both numerical and visual outputs for the analysis of adhesion structures and colocalization of stained proteins in confocal microscopy images. It supports fully automated workflows while allowing manual corrections – such as structure editing and parameter adjustment – at key stages of processing, segmentation, and classification. Results are automatically exported in spreadsheet format for further analysis.

Offered as open-source software, the tool is accessible and adaptable, positioning it as a viable alternative to existing platforms. Planned improvements include enhanced automation (e.g., label merging, multi-cell detection), user interface refinements, and support for custom parameter presets to streamline future analyses.

## 7. Authors’ Contribution

**Joanna Hajduk:** Writing – original draft, Investigation, Validation, Visualisation. **Patrycja Twardawa:** Writing – original draft, Data curation, Methodology, Software. **Zenon Rajfur:** Writing – review & editing, Supervision.

## 9. Acknowledgment

This study was funded by the ‘Research Support Module’ (RSM/89/HJ and RSM/94/LO) as part of the ‘Excellence Initiative – Research University’ program at the Jagiellonian University in Kraków.

