## Supplementary materials for "Software for semi-automatic analysis of microscopic images of adhesion structures and protein colocalization in cells"

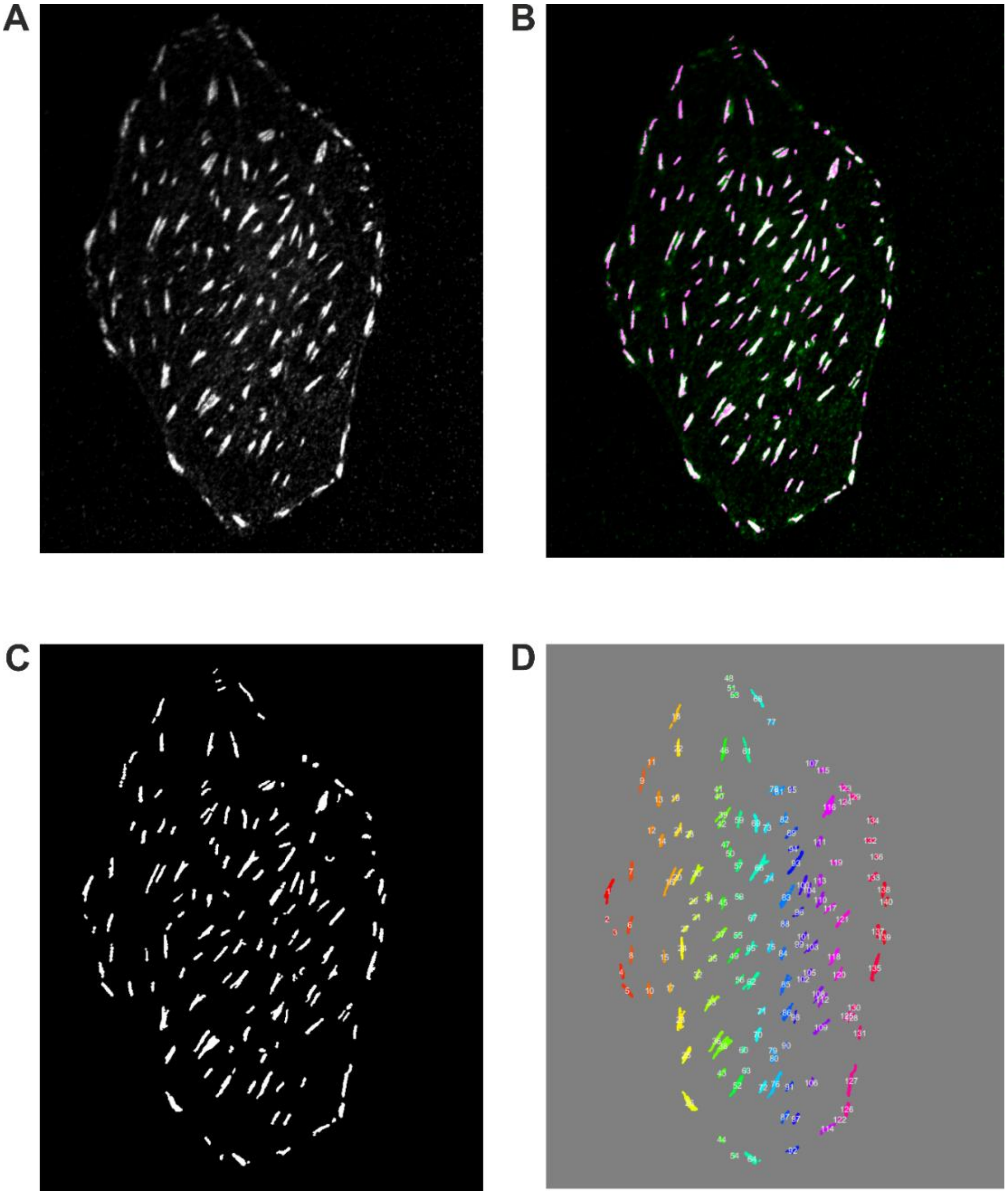


**Fig S1.** Exemplary result of segmentation of adhesion structures in U2OS cell seeded on glass covered with fibronectin. **A)** Original image of stained Talin-1 in U2OS cell with adjusted contrast. **B)** Overlay of found structures (purple / white) on the original image (green). **C)** Binary mask of found adhesion structures. **D)** Found adhesion structures with labels. Label colors are assigned randomly.

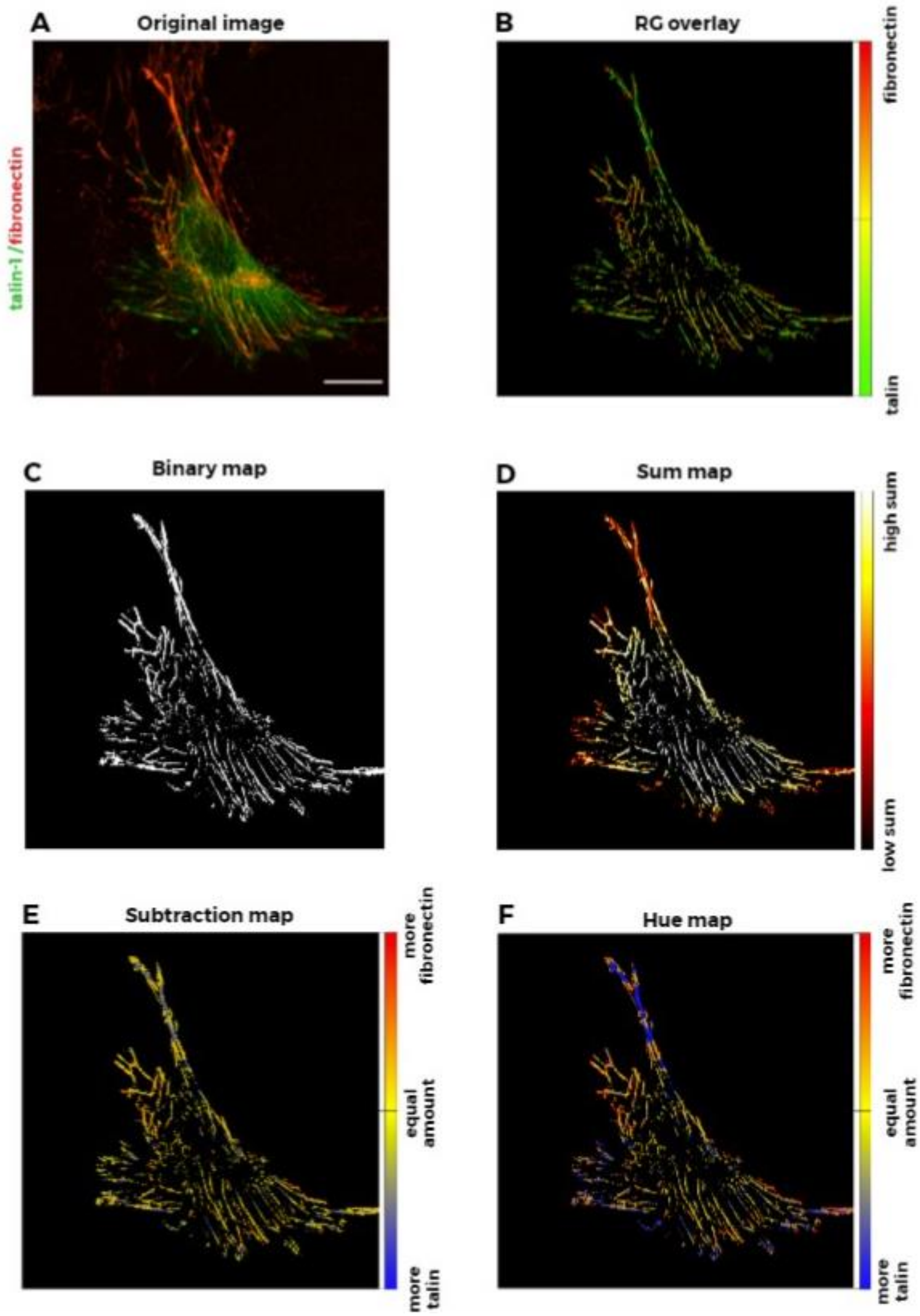


**Fig. S2.** Stages of colocalization map preparation with exemplary results of two image channels. The image shows a NIH/3t3 cell stained for Talin-1 (green channel) and fibronectin (red channel). Scale bar: 20 µm. **A)** Original image with adjusted contrast. **B)** Overlay of both channels after segmentation of cellular structures. **C)** Binary mask of cellular structures detected in both image channels. **D)** Sum map of normalized red and green channel intensities within the segmented regions. **E)** Subtraction map of normalized red and green channel intensities, indicating the relative prevalence of each component. **F)** Hue map representing which channel dominates in each masked region, independent of object intensity.

### USER’S MANUAL

#### for the *Focal Adhesions* and *Colocalization* Software

##### Overview

This software was designed specifically for the semi-automatic analysis of adhesion structures in microscopy images. Program also analyzes colocalization between two fluorescent channels, providing both visual and quantitative assessment. This software tool automates essential workflow, such as ROI selection, contrast adjustment, denoising, background subtraction, filtering, segmentation etc., while still preserving the ability for users to intervene manually when needed.

One of the script add-ons focuses on the segmentation and analysis of cellular adhesion structures, providing quantitative measurements for each detected object. Multiple visualization modes and labeled maps are available to facilitate visual inspection and validation of identified adhesion structures. For double-stained images, the software offers advanced colocalization features, including correlation coefficient calculation (Pearson’s, Spearman’s) between selected channels and generation of various maps (sum, subtraction, channel overlay on the binary structure mask, and hue maps), enabling detailed assessment of emission intensity and spatial relationships between stained structures. This tool is particularly useful for researchers investigating structure-specific properties and interactions between adhesion-related components across a range of imaging conditions.

##### Prerequisites

- MATLAB (Version R2021a or later), with required toolboxes:
    - Image Processing Toolbox;
    - Statistics and Machine Learning Toolbox.

##### Workflow instruction

The script workflow and the proposed order of actions are presented in the main body of the publication (Fig. 1). Users should refer to the block diagram for guidance on the sequence of script sections. A detailed description, including suggestions for parameter adjustments, recommended value ranges, and potential sources of errors, is provided in the section below. For cases not covered in this manual, users are encouraged to contact the authors.

##### General remarks

In case of troubleshooting, warnings encountered during execution will not stop the script but may indicate incomplete data loading or partial processing at a specific stage. Users are advised to pay close attention to console prompts, as the appearance of a warning may carry important information relevant to the overall analysis.

If an error occurs, the script cannot continue operating correctly. Users should follow the instructions provided in the error message and ensure that the underlying issue is resolved – either by correcting the input data or addressing the cause of the error – before proceeding to the next steps or sections of the script.

##### Manual legend

**WARNING:** Sections labeled with ‘WARNING’ contain important information related to the operation of the corresponding script section. It is strongly recommended to read the content of these messages carefully.

**TROUBLESHOOTING:** Sections labeled 'TROUBLESHOOTING' contain tables of error and warning descriptions along with proposed solutions. If an unlisted error occurs or the suggested solution does not yield satisfactory results, users are encouraged to contact the authors of this work.

**NOTE:** Sections labeled 'NOTE' provide suggestions intended to support the user's workflow or enhance understanding of the script's behavior.

**STEP 1: Initialize parameters, choose add-ons and adjustable variables**

1. Run 'VARIABLE INITIALIZATION / RESET' to initialize non-adjustable parameters.
2. Run the 'ADD-ON CHOICE (USER INPUT)' section to select the desired add-ons (focal adhesions feature, colocalization feature, or both). Add-ons are enabled by setting their corresponding variables to 1 and disabled by setting them to 0.
3. To modify user-adjustable variables, navigate to the 'OPTIONS (USER INPUT)' section, update the desired values and run the section. All adjustable parameters are detailed in Tab. S1.

   **WARNING:** In the sections between the clauses "*adjustable parameters*" and "*end of section for modification / user input*", all adjustable parameters, flags, and method selection options are explicitly detailed to allow users to modify script behavior. Users are advised to stay within the specified ranges to ensure correct script operation.

   **NOTE:** This section may be updated and reloaded at any point during script execution, particularly if users need to reprocess or reclassify images to refine the performance. In such cases, all necessary values should be adjusted, and the section reloaded, after which users can return to the sections where they left off in the script and continue without data loss.

**STEP 2: Upload image channel file/-s**

1. Run the 'IMAGE DATA READ & DATA PREPARATION' section to load the image data (in '.tiff' format). If only the focal adhesions add-on is used, a single image channel file should be loaded. If the colocalization add-on is enabled, two selected image channel files are required. During the image loading process, pixel size information will be extracted from the file metadata. If pixel size data cannot be retrieved or is not available, the user will be prompted via the console to input the value manually. During this stage, also the diameter for ring areas will be counted, based on the pixel size data.

**TROUBLESHOOTING:**

| Number of error/warning | Error / warning prompt | Error / warning description | Recommended solution |
|---|---|---|---|
| E1 | ERROR: File read error for COLOCALIZATION add-on. Check the format and size of files (TIFF). | The error indicates a failed image read in the *Colocalization* add-on. | Please check the file format and ensure that the image file is not corrupted. |
| E2 | ERROR: File read error for FA add-on. Check the format and size of files (TIFF). | The error indicates a failed image read in the *Focal Adhesions* add-on. | |
| E3 | ERROR: File read error. Check root file and try again. | This error may appear alongside the E1 / E2 prompts and indicates that the file cannot be read. | |
| E4 | ERROR: Ring diameter could not be obtained. Please check image metadata / provide information about pixel size in | This error occurs when the conversion from real-world units to pixel units fails. | Please check the variable *ring_diameter_µm* in the *ADJUSTABLE PARAMETER VALUES: OPTIONS (USER INPUT)* |

| | | | |
|---|---|---|---|
| | real units. You must repeat image read. | | section. Ensure that the *ring_diameter_µm* value is numeric, non-empty, and greater than 0, and that it does not exceed the shorter dimension of the image. The value should be provided in µm.<br>Also, verify that the pixel size has been correctly determined. If it could not be extracted from the image file metadata, the script prompts the user to enter it manually via the console. If this prompt was skipped or left blank, the pixel size value may be missing. In such cases, re-run the image reading section and provide the pixel size when prompted. |
| W1 | WARNING: Unexpected pixel size. Check file for metadata corruption. Add size of pixel in µm: | This warning indicates that the pixel size should be verified by the user, as the file metadata may be corrupted or the pixel size information could not be retrieved properly. | When prompted, provide the pixel size in micrometers (µm). Do not skip the prompt or leave the input blank or non-numeric. If this issue occurs, repeat the image reading section and ensure the correct value is entered. |
| W2 | WARNING: No data about pixel size in root files. Add size of pixel in µm: | This warning indicates that the file metadata was read successfully and is not corrupted, but it does not contain information about the pixel size. | |
| W3 | WARNING: Information about pixel size has already been provided. You can proceed with distance recounting. | This warning indicates that pixel size data already exists in script workspace. | No action is required. This warning indicates that a user action was blocked to prevent unintended script behavior. |
| W4 | WARNING: You have already recounted the data to real distance units. Operation cannot be repeated. | This warning indicates that pixel size data already exists, and a double unit conversion has been prevented to ensure data consistency. | |

**NOTE:** It is advisable to use consistent contrast enhancement settings across all images in a larger batch, as changes in contrast directly affect the visualized object intensities and will influence the final results of emission intensity analysis. Contrast adjustments are applied after value normalization and initial denoising, and are scaled proportionally to the actual image maximum prior to the full image processing stage.

**NOTE:** Please note that the *ring area* diameter should be specified by the user in micrometers (µm), and the corresponding pixel distance will be calculated based on the provided real-world measurement.

**STEP 3: Set contrast settings**

1. Users can adjust contrast settings for visualization during later stages of script execution. Since original images often exhibit low brightness and contrast, adjusting these settings can improve usability. The script allows the use of either automated contrast settings or user-defined values for image analysis. These options, along with contrast limits, can be modified in the ‘adjustable parameters’ section. It is recommended to use the parameter ranges provided in Table S1.

**STEP 4: *CELL ROI* selection and modification**

**Automatic *CELL ROI* selection**

1. Run 'AUTO CHOICE OF ROI' section for automatic ROI selection.
2. If multiple objects are detected (as shown in the image below), select one by typing its corresponding number in the console when prompted. You can enable automatic selection of the main cell body – defined as the largest visible area in the image – by setting the *main_area_flag* variable in the "*adjustable parameters*" section. If *main_area_flag* is set to 1, the largest area will be selected automatically. If set to 0, the user will be prompted to manually choose the desired cell area.

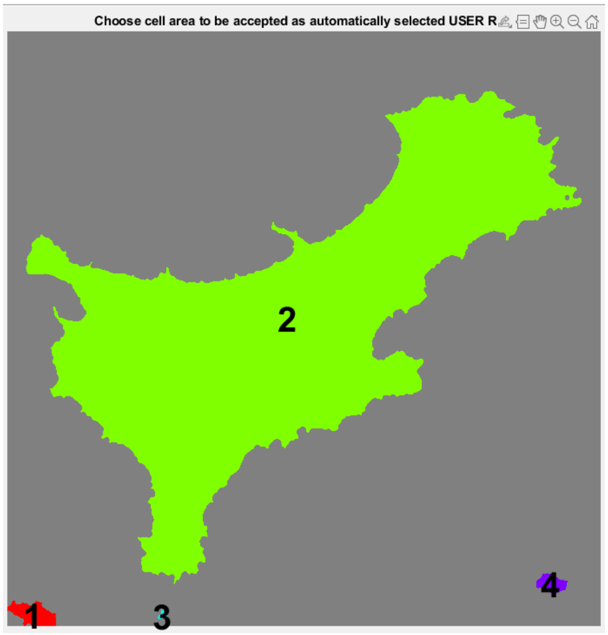


*CELL ROI selection with manual main cell body selection enabled (main_area_flag = 0).*

3. A preview will be generated to allow the user to view the ROI in two displayed images: a binary map of the extracted USER ROI and an overlay of the USER ROI on the original image.

**NOTE:** If multiple cells are present in the microscopy image, it is advisable to select the main cell body manually – by setting the *main_area_flag* value to 0 – to allow switching between cells, rather than defaulting to the largest one for analysis.

**Manual *CELL ROI* selection and modification**

If the user is not satisfied with the automated *CELL ROI* selection, or prefers manual selection, the *CELL ROI* can be either completely replaced or manually adjusted.

**Manual *CELL ROI* selection**

1. To manually outline the CELL ROI, run the 'USER ROI CHOICE: MANUAL CHOICE OF CELL ROI' section. Note that initiating a new manual selection will overwrite any previously saved results. If only an adjustment to the currently selected CELL ROI is needed, use the sections located below the 'CELL ROI MODIFICATION: MANUAL CORRECTION OF ROI' clause.
2. Double-click to close the selection envelope around the chosen CELL ROI area.
3. Press any key to return to the script workspace and console, and to confirm the selection.

**Manual *CELL NUCLEUS ROI* selection**

1. To outline the *CELL NUCLEUS ROI*, run 'USER ROI CHOICE: MANUAL CHOICE OF CELL NUCLEUS ROI' section. An example of manual *cell nucleus ROI* selection is shown in the image below.
2. Double-click to close the selection envelope around the chosen *cell nucleus ROI* area.
3. Press any key to return to the script workspace and console, and to confirm the selection.

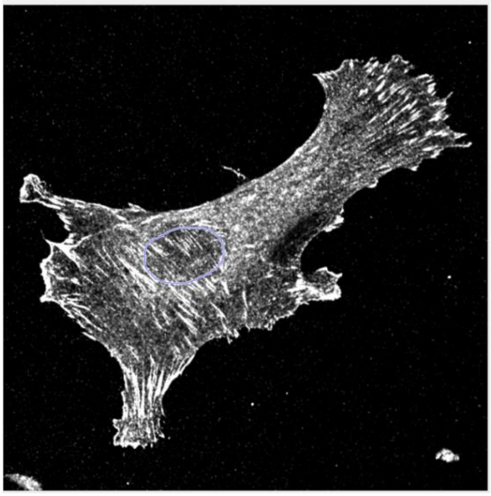

*Example of manual CELL NUCLEUS ROI selection during the outlining process.*

**WARNING:** Selection of the *cell nucleus ROI* must be performed manually by the user and will not be executed if the script is run in fully automatic mode. Modification or reversion of the selection is not implemented for the *cell nucleus ROI*. The selection can be performed multiple times; each new selection will override the previous binary map.

**Manual CELL ROI modification**

1. To modify an existing *CELL ROI*, either:
   - Use the 'DELETE OBJECTS FROM CELL ROI' section to remove objects from the *cell ROI*
   - Use the 'ADD OBJECTS TO CELL ROI' section to add objects to the *cell ROI*
2. Double-click to close the selection envelope around the modified area. The results of the user intervention will be displayed immediately in the display window.
3. Press any key to return to the script workspace and console, confirming the selection.
4. After any manual modification, the image can be reverted to its previous state using the 'UNDO /REDO CELL ROI MANUAL CHANGES' section of the script, which supports one level of undo and redo actions.

**NOTE:** The user can make multiple selections during a single session of deleting or adding objects. All objects added to or removed from the *CELL ROI* during one marking session will be treated as a single modification, and the entire set of changes will be subjected to 'UNDO / REDO' operations.

#### TROUBLESHOOTING

| Number of error/warning | Error / warning prompt | Error / warning description | Recommended solution |
|---|---|---|---|
| E5 | ERROR: No add-on selected. Choose either FA, COLOCALIZATION, or | This error appears when no add-on has been selected. | Choose either the *Focal Adhesions* or *Colocalization* add-on by setting the corresponding variable value to |

| | | | |
|---|---|---|---|
| | both. Check USER INPUT variables. | | 1 in the *ADD-ON CHOICE (USER INPUT)* section of the script. |
| E6 | ERROR: Image is too small for automated ROI correction. Try to perform CELL ROI choice manually. | The small image size prevents the automatic *CELL ROI* selection algorithm from operating properly. | It is advisable to verify that the correct image has been provided for analysis. If the data has been confirmed and the error still occurs, proceed with *manual CELL ROI* selection. |
| E7 | ERROR: Chosen channel for ROI extraction is not read due to USER'S choice. Please check USER INPUT variable section ('ROI_main_ch' variable). | The main channel from which the *CELL ROI* should be extracted for the *Focal adhesions* add-on (when both *Focal Adhesions* and *Colocalization* add-ons are used) has been specified outside the available range. For example, the user may have selected the R and G channels, but the B channel was provided for *CELL ROI* selection. | Select one of the available channels by modifying the *ROI_main_ch* variable in the *ADJUSTABLE PARAMETER VALUES: OPTIONS (USER INPUT)* section. The value should correspond to the number of one of the provided channels, as specified in the *ch_coloc* variable, where the channel labels and their corresponding numbers are as follows: 1 – R channel, 2 – G channel, 3 – B channel. |
| E8 | ERROR: Chosen channel for UNDO / REDO operations is not read due to USER'S choice. Please check USER INPUT variable section ('ROI_main_ch' variable). | | |
| E9 | ERROR: No object found. Automated CELL ROI extraction failed. Please, choose CELL ROI manually and check image data beforehand. | No object was found in the result of the algorithm's operation. | Proceed with the manual *CELL ROI* choice instead of automatic selection. If the error appears consistently with your data, users are encouraged to contact the authors of this work |
| E10 | ERROR: Failed attempt of CELL ROI choice. Check data and try again before proceeding further. | Manual CELL ROI selection has failed. | The user should ensure that the selection window was not closed without making a selection via a double-click of the left mouse button (e.g. by exiting the window or pressing any key). Please repeat the operation. If the issue persists, check the input data. |
| W5 | WARNING: Your chosen CELL ROI area could not be accepted. The biggest area was chosen instead. | The selected object number does not correspond to any of the detected objects. | No action is required. This warning indicates that a user action was blocked to prevent unintended script behavior. The largest object will be selected automatically. If a different object needs to be chosen, the procedure must be repeated. |
| W6 | FOCAL ADHESIONS ADD-ON \| WARNING: UNDO / REDO operation could not be performed. No changes have been applied. | The UNDO / REDO operation could not be applied due to an internal processing error. | Check the validity of the input variables, especially *Iout_ncirc, temp_Iout_ncirc, Iout_ncirc_old, and temp_Iout_ncirc_old*. Note that if no add or delete operations have been performed, the ‘UNDO / REDO’ section cannot proceed due to the lack of data – this is expected behavior. |

#### STEP 5: Image Processing and Segmentation

During the first stage, image processing is performed to enhance segmentation for the purposes of the *Focal Adhesions* and *Colocalization* add-ons. The processing steps differ between the two add-ons, as they employ distinct approaches. In the second stage, binary segmentation maps are generated. The result of the segmentation – highlighting the cellular structures within the *CELL ROI* – is referred to as the *SEGMENTATION ROI*.

1. Execute the 'IMAGE PROCESSING, SEGMENTATION' section to:
   a. Apply denoising algorithms
   b. Perform background subtraction
   c. Apply image filtering using the selected parameters (including fibrillar structure enhancement for the *Focal Adhesions* add-on)
   d. Perform segmentation using the chosen parameters
2. Review the segmentation results for both the whole cell and the cell nucleus in the display windows. If the results are unsatisfactory, adjust the image processing and/or segmentation parameters by modifying the variable values in the 'OPTIONS (USER INPUT)' section. Descriptions of the variables can be found in the script documentation and in Table S1.

**NOTE:** Denoising can be performed using either a pre-trained *DnCNN* network or morphological operations. While *DnCNN*-based denoising substantially improves image quality, it may require significant computational resources. Therefore, this method is not recommended for use on older or low-performance devices.

**NOTE:** The image channel used for segmentation should be specified in advance, but can be changed at any time before segmentation by modifying the values in the 'OPTIONS (USER INPUT)' section *(ch_tal* variable for the *Focal Adhesions* add-on and *ROI_main_ch* variable for the *Colocalization* add-on). The expected input for these variables is the channel number: R = 1, G = 2, and B = 3.

**TROUBLESHOOTING**

| Number of error/warning | Error / warning prompt | Error / warning description | Recommended solution |
|---|---|---|---|
| E5 | ERROR: No add-on selected. Choose either FA, COLOCALIZATION, or both. Check USER INPUT variables. | This error appears when no add-on has been selected. | Choose either the *Focal Adhesions* or *Colocalization* add-on by setting the corresponding variable value to 1 in the *ADD-ON CHOICE (USER INPUT)* section of the script. |
| E11 | ERROR: Segmentation failed. Check workspace data and try again after repeating former steps. | This error indicates a segmentation failure, resulting in the absence of object assignment. | It is advised to check the workspace data (e.g. to ensure that the input image contains visible structures) and the post-processed image, particularly to verify that it has not been overprocessed to the point where the structures are no longer sufficiently visible for the segmentation algorithms. The user can inspect either the *G_tal* or *G_col* image (depending on the add-ons used – *Focal Adhesions* or *Colocalization*, respectively) by entering the command 'imshow(G_tal);' or 'imshow(G_col);' in the console window. |
| E12 | ERROR: Error during image processing and segmentation. Objects have not been designated. Please check data. | No objects were found in the image, and the binary map could not be generated. | |
| W7 | WARNING: Denoising failed. You can proceed further without image modification. | The error indicates that denoising procedure failed. The script will proceed without applying denoising. | Denoising failure may result from using the *DnCNN*-based approach, which requires significant computational resources. To use morphological-based denoising |

| | | | |
|---|---|---|---|
| | | | instead, set the *denoise_flag* value to 2. If both procedures fail, ensure that the original input image exists in the workspace and is valid. |
| W8 | WARNING: Background subtraction failed. You can proceed further with unsubtracted background. | This error indicates that the background subtraction procedure failed. The script will proceed without applying background subtraction. | Check the validity of input parameters and workspace data (e.g. to ensure that the data has not been accidentally cleared). If the data is correct, try adjusting the size of the structural element for background subtraction (*se* variable) by selecting a smaller radius. Also, ensure that the input data is in the correct format for each variable. |

#### COLOCALIZATION ADD-ON

The functions described below are specific to the Colocalization add-on and correspond to script sections that should be used only if the user intends to utilize the features of this add-on.

##### STEP 6C: Colocalization analysis

1. Run the 'COLOCALIZATION ADD-ON | MAP PREPARATION & SEGMENTATION' section to:
    a. Generate the initial *Sum Map* for the whole cell and nucleus regions
    b. Perform segmentation using the selected method and generate binary segmentation masks for whole the cell and nucleus regions
    c. Generate the *Sum Map*, *RGB Overlay Map*, *Subtraction Map*, and *Hue Map* for the segmented cellular structures in the whole cell and nucleus regions

**NOTE:** All generated images and charts are automatically saved in both FIG and TIFF formats in the same directory as the original input image. Datasets related to correlation coefficients are saved in XLSX spreadsheet format in the same directory.

##### TROUBLESHOOTING

| Number of error/warning | Error / warning prompt | Error / warning description | Recommended solution |
|---|---|---|---|
| E13 | COLOCALIZATION add-on \| ERROR: Map preparation error. Please check before continuation. | The colocalization display maps could not be generated due to errors during file reading, processing, or saving. | Check the validity of input parameters and workspace data (e.g. to ensure that the data has not been accidentally cleared). Verify the save path length and validity by checking the *name2CH* and *filepath2CH* variables. Also, ensure that the binary maps generated during segmentation for the Colocalization add-on were created correctly and that the objects are visible. You can use the commands 'imshow(B6col);' and 'imshow(B6col_nucl);' via console window to inspect the results. |

##### STEP 7C: Correlation coefficient calculation

1. Run the 'COLOCALIZATION ADD-ON | CORRELATION COEFFICIENT CALCULATION' section to compute Pearson's and Spearman's correlation coefficients for colocalization analysis using the two provided image channels. Values are calculated separately for the whole cell and the nucleus.

**TROUBLESHOOTING**

| Number of error/warning | Error / warning prompt | Error / warning description | Recommended solution |
|---|---|---|---|
| E14 | COLOCALIZATION add-on \| ERROR: Correlation coefficient calculation failed. Please check data and try again. | The correlation coefficients could not be properly calculated or saved. | Check the validity of input parameters and workspace data (e.g. to ensure that the data has not been accidentally cleared). Verify the save path length and validity by checking the *name2CH* and *filepath2CH* variables. |

##### STEP 8C: Data display and saving

The results are automatically displayed and saved in an XLSX format spreadsheet during the execution of *STEP 7C*. The spreadsheet is saved in the folder containing the first image channel used for the colocalization add-on.

#### FOCAL ADHESIONS ADD-ON

The functions outlined below are specific to the *Focal Adhesions* add-on and correspond to script sections intended solely for users who wish to apply the features of this module. Complete this section if you have set the add-on flag to 1 (*FA* variable).

##### STEP 6FA: Analysis of emission intensity within concentric bands of a specified diameter ('*ring areas*')

1. Run the 'FOCAL ADHESIONS ADD-ON | COUNTING INTENSITY IN THE BANDS OF GIVEN DIAMETER' section to calculate intensity within concentric rings (so-called '*ring areas*'). The diameter of these rings is specified by the user via the *ring_diameter_um* variable in the 'OPTIONS (USER INPUT)' section. The selected diameter is then converted to pixel units. The user can also choose whether holes within the cell body should be included in the '*ring areas*', or whether the rings should overlap each hole (controlled by the *ring_hole_flag* variable: set to 1 – holes will be enveloped; set to 0 – holes will be ignored).
2. To optionally visualize a specific '*ring area*', run 'FOCAL ADHESIONS ADD-ON | USER CHOICE: DISPLAY CHOSEN RING AREA'. By default, the first ring is displayed. To visualize a different '*ring area*', modify the *ring_number* variable.

**TROUBLESHOOTING**

| Number of error/warning | Error / warning prompt | Error / warning description | Recommended solution |
|---|---|---|---|
| E15 | FOCAL ADHESIONS ADD-ON \| ERROR: Objects in chosen rings of given diameter cannot be counted. Check data and try again. | An error occurred during the estimation of consecutive ring areas. | Please check the data, especially the ring diameter (ensure that the *ring_diameter_um* value does not exceed the cell size limits; start with smaller values), and verify the presence of a binary map (i.e., a |

| | | | |
|---|---|---|---|
| | | | non-empty *segmentation ROI*). Then try again. Consider changing the value of the *ring_hole_flag* variable to determine whether empty spaces within the cell body are causing the counting malfunction. |
| E16 | FOCAL ADHESIONS ADD-ON \| ERROR: There is no object to erode from. Check data and assign ROI again. | The binary image of cell contains no found object. | Consider adjusting the segmentation parameters (by reloading the section containing adjustable settings) to reveal the objects in *segmentation ROI*, or add them manually if segmentation fails, in order to proceed with this stage. |
| E17 | FOCAL ADHESIONS ADD-ON \| ERROR: Ring diameter not provided. Please check your data and try again. | The variable containing the ring diameter is either empty, has been cleared, or contains non-numeric data. | Provide a numeric value for the *ring_diameter_um* variable. Ensure that the value does not exceed the cell size limits – try starting with small values. |
| E18 | FOCAL ADHESIONS ADD-ON \| ERROR: Failure during processing of ring areas. Check data and assign ROI again. | A general error indicating that the data provided as the function entry prohibits it from operating correctly. | Check whether the input data is correct – for example, whether the binary map for the *segmentation ROI* exists and whether numeric values have been provided for *ring_hole_flag* and *ring_diameter_um*. Ensure that *ring_hole_flag* is set to either 0 or 1. Verify also that the value of *ring_diameter_um* does not exceed the size of the cell (i.e., that the band is not larger than the cell itself) and that it has been correctly provided in µm unit. |
| W9 | FOCAL ADHESIONS ADD-ON \| WARNING: Ring area number exceeds the number of found ring areas. The maximum ring area number is … . Reload data and try again. | The '*ring area*' of the selected number cannot be displayed due to an invalid ring number value provided by the user. | Check whether the ring number has been provided correctly (via the *ring_number* variable), and refer to the console prompt to determine the maximum allowed value. Also verify that the variable contains numeric data. |
| W10 | FOCAL ADHESIONS ADD-ON \| WARNING: Ring area image cannot be displayed. Check data. | The 'ring area' cannot be displayed. | Check the validity of the input data. Ensure that the original image and the structure containing ring data are not empty or deleted. If necessary or if the structure containing the ring data (*ir_rings* variable) is empty, re-run the 'FOCAL ADHESIONS ADD-ON \| COUNTING INTENSITY IN THE BANDS OF GIVEN DIAMETER' section. |

**STEP 7FA: Object selection and parameter estimation (initial / I)**

1. Run 'FOCAL ADHESIONS ADD-ON | OBJECT SELECTION & PARAMETER ESTIMATION (1)' section to:

a. Calculate the object properties based on the previously estimated *segmentation ROI*. The properties include: area, centroid, circularity, perimeter, major axis length, minor axis length, max/min dimension ratio, orientation, intensity centroid, mean intensity, minimum intensity, and maximum intensity.
b. Assign labels to all found objects.
c. Perform object classification based on a combination of properties such as shape, area, and intensity level, optionally including or excluding objects manually added by the user (as specified by the *manualcorr_flag* variable). This step generates binary and label maps of the objects identified as focal adhesions, along with a table structure containing parameter data for all objects (*temp* variable).
d. Shift the labels to reflect the classification results, if any objects were rejected based on the classification criteria.
e. Identify the objects located over the cell nucleus based on the updated *segmentation ROI* after classification (binary map of focal adhesions) and the *cell nucleus ROI*. These objects will be added to the structure containing data for all detected objects (*temp* variable).
f. Review the results using the displayed label and binary maps (an example binary map of the detected objects is shown in the image below).

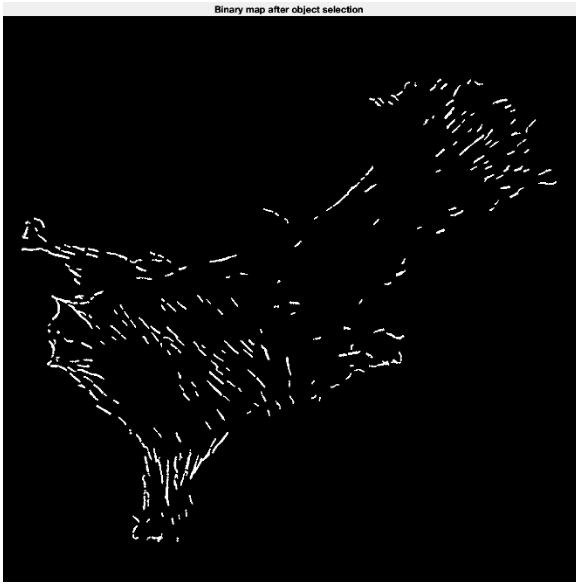


*Example of a binary map showing the detected focal adhesions, based on the segmentation ROI after the classification stage.*

#### TROUBLESHOOTING

| Number of error/warning | Error / warning prompt | Error / warning description | Recommended solution |
|---|---|---|---|
| E19 | FOCAL ADHESIONS ADD-ON \| ERROR: Labelling failed. Check data and try again. | The error indicates a general parameter estimation / classification / labelling failure. | Check for any errors displayed in the console prompt alongside this message. Verify the validity of the input data and re-run the previous script stages if necessary. Ensure that objects have been detected within the *segmentation ROI* after segmentation. If needed, improve segmentation performance by adjusting the variables in the *adjustable parameters* section, or add objects manually if automatic segmentation continues to fail. |

| | | | |
|---|---|---|---|
| E20 | FOCAL ADHESIONS ADD-ON \| ERROR: Labelling failed. Object selection failed. Check data and try again. | A failure occurred during object classification or labelling, following parameter estimation. | Check the validity of the object data (*temp* variable) while running the script step by step. Verify the data passed to the *talclassify* function, which is used within the *talCountParams* method. If necessary, adjust the parameters and re-run the segmentation stage after modifying the segmentation settings (via the *adjustable parameters* section), or manually add or delete objects from the binary maps before returning to this section. |
| W11 | FOCAL ADHESIONS ADD-ON \| WARNING: Processing flag turned OFF. | Processing was disabled due to missing data. | Please check whether the previous script stages have already been completed, and repeat any necessary steps (e.g., segmentation). |
| W12 | FOCAL ADHESIONS ADD-ON \| WARNING: Classification finished. No object has been rejected based on provided criteria. | No objects have been rejected based on the active classifier criteria. | This warning is not an error, but rather an indication of specific classifier behavior, intended to alert the user that all objects meet the classification criteria. If this outcome is not intended, the user can modify the classification parameters in the *adjustable parameters* section ('OPTIONS (USER INPUT)', under the 'FOCAL ADHESIONS ADD-ON' and 'SEGMENTATION & CLASSIFICATION PARAMETERS' clauses). |
| W13 | FOCAL ADHESIONS ADD-ON \| WARNING: Object status & label maps based on localization relative to nucleus were not prepared. | Objects above the cell nucleus were not successfully detected. This error does not occur if no such objects are present and the relevant variables and *cell nucleus ROI* have been correctly specified. | Check whether the *cell nucleus ROI* has been selected manually. The *binaryImage2* variable should not be empty and must contain the user-defined *cell nucleus ROI*. You can use the '*imshow(binaryImage2);*' command in the console to verify the ROI. Step through the *talCountParams* method and verify the validity of the input data passed to the *talFindAboveNucleus* method, which is called within *talCountParams*. |

**NOTE:** Objects are excluded based on the following criteria: low overall intensity combined with small area, low emission intensity in larger objects, or small area combined with circular shape and low emission intensity. The user should note that the area thresholds for circular and non-circular objects may differ (*object_area_thresh_circ_itsty* and *object_area_thresh_itsty* variables in the *adjustable parameters* section). For optimal classifier performance when detecting focal adhesions, it is recommended to set the area threshold for circular objects higher than for fibrillar ones. The user should also check the resulting image for undetected nascent adhesions, which may appear as small circular structures. If automatic adjustment does not yield a balanced segmentation outcome, the user can always switch to manual mode to mark any missing focal or nascent adhesion structures.

**STEP 8FA: Classified *segmentation ROI* modification – adding & removing objects**

1. To modify the *segmentation ROI* after segmentation and initial classification, run one of the following:
    a. 'FOCAL ADHESIONS ADD-ON | DELETE OBJECTS FROM SEGMENTATION ROI' – to remove objects from the *segmentation ROI* after classification.
    b. 'FOCAL ADHESIONS ADD-ON | ADD OBJECTS TO SEGMENTATION ROI' – to add objects to the segmentation ROI after classification.
    c. 'FOCAL ADHESIONS ADD-ON | UNDO / REDO FOR SEGMENTATION ROI' – to revert the last change.

**NOTE:** The user can add or delete multiple objects during a single addition or deletion session; this will be recognized as a single action for the purpose of the 'UNDO / REDO' option. The action is finalized by pressing any key, as described in the section instructions and indicated in the console prompt.

**NOTE:** Before proceeding further, the user must verify whether the *segmentation ROI* map is satisfactory, as this is the final stage where modifications to its structure can be made. If changes are required after executing subsequent stages – such as label merging – the user must return to the earlier sections of the code responsible for modifying the *segmentation ROI*. This will result in the loss of all results generated in the later stages.

**TROUBLESHOOTING:** The possible errors are the same as those that may occur during *cell ROI* modification. Refer to the descriptions of errors E5, E7, and E8, and warnings W5 and W6 in the *TROUBLESHOOTING* subsection of the *STEP 4: CELL ROI Selection and Modification* section of the manual.

**STEP 9FA: Object selection and parameter estimation (final / II)**

1. After making all necessary modifications, run 'FOCAL ADHESIONS ADD-ON | OBJECT SELECTION & PARAMETER ESTIMATION (2)' to update the parameters:
    a. Calculate the object properties based on the *segmentation ROI* after the initial classification stage.
    b. Assign new labels to all detected objects if the object map has changed due to manual modifications made by the user.
    c. Select the objects marked by the user as potentially disposable by the classifier (by modifying the *manualcorr_flag* variable) if they do not meet the classification criteria, or as non-disposable if the user has chosen to override the classifier rules for manually added objects. The classification rules will still apply to all other objects detected automatically and can only be overridden by adjusting the parameters used by the classifier, via the adjustable parameters section ('OPTIONS (USER INPUT)' section; variables listed under 'FOCAL ADHESIONS ADD-ON' and 'SEGMENTATION & CLASSIFICATION PARAMETERS' clauses.
    d. Perform object classification based on a combination of properties such as shape, area, and intensity level, optionally including or excluding objects manually added by the user (as specified by the *manualcorr_flag* variable). This step generates binary and label maps of the objects identified as focal adhesions, along with a table structure (*temp* variable) containing parameter data for all objects.

e. Shift the labels to reflect the classification results, if any objects were rejected based on the classification criteria.
f. Review the results using the displayed label and binary maps.

**NOTE:** The user should be aware that after this stage, modifications to *segmentation ROI* will no longer affect the label map. To implement such changes in subsequent results, it will be necessary to return to the earlier stages of the script.

**TROUBLESHOOTING:** The possible errors are the same as those that may occur during initial object selection and parameter estimation. Refer to the descriptions of errors E19, E29, and warnings W11-W13 in the *TROUBLESHOOTING* subsection of the *STEP 7FA: Object selection and parameter estimation (initial / I)* section of the manual.

##### STEP 10FA: Labels modification

1. Use the 'FOCAL ADHESIONS ADD-ON | CONNECTING LABELS / MERGING OBJECTS' section to merge selected labels. The label merging process is similar to the ROI modification stage:
   a. the label map is displayed, allowing the user to select objects whose labels are to be merged into a single structure. Label merging is performed by drawing a selection area on the screen – any object even partially enclosed within this area will be included in the merged structure.
   b. After confirming the selection (by pressing any key or double-clicking the left mouse button), the label numbers will be automatically reassigned after each change.

**WARNING:** The label merging process cannot be undone. Please proceed with caution, or exit the selection window before confirming the selection by pressing any key or double-clicking the left mouse button.

**NOTE:** While labels can be merged, they cannot be split at this stage. If users wish to divide objects into smaller structures manually, such modifications must be made directly on the segmentation ROI before the classification stage. Classification and labeling must then be repeated, and any previously performed merging will be lost.

**NOTE 2:** Note that multiple selections can be made during a single session. If those operations slow down your computer, close the window and wait for the console message: '*Label merging finished. Object status & label maps based on localization relative to the nucleus have been successfully generated*', before proceeding with next selections.

**NOTE 3:** Merged objects will not be connected by any artificial additions – not even a thin line – in order to preserve the original segmentation conditions. The user can influence object shape only during the ROI modification stages.

##### TROUBLESHOOTING

| Number of error/warning | Error / warning prompt | Error / warning description | Recommended solution |
|---|---|---|---|
| E21 | FOCAL ADHESIONS ADD-ON \| ERORR: Ratio property of objects was not reassigned properly. Check data and try again. | The ratio property (the ratio between the object's major and minor axis lengths) cannot be accurately transferred to the structure after label merging. | Check the validity of the input data for the *talRatioGoI* method. Ensure that the parameters table (temp variable) exists and contains detected objects. Try repeating STEP 9FA (final object selection and parameter estimation), and then proceed with label merging again. Note that in this case, any previous label merging results will be lost. |

| | | | |
|---|---|---|---|
| W14 | FOCAL ADHESIONS ADD-ON \| WARNING: Only one object selected. Label merging not performed, try again. | Only one object was selected during the label merging process. Labels cannot be merged unless at least two objects are selected. | Run the section again and pay close attention to the selection marking. Note that objects must be at least partially enclosed within the selection area to be included in the label merge into a single structure. Confirm the selection by pressing any key or by double-clicking the left mouse button. |
| W15 | FOCAL ADHESIONS ADD-ON \| WARNING: Labeled objects not merged. Previous labels valid. Check data or proceed. | This warning indicates a malfunction in the relabeling process. The user can proceed with other sections of the script, as the previous data has been preserved; however, it is recommended to verify the data before continuing. | A malfunction in the relabeling process may result from interrupting the software during recalculation or from accidental modification of workspace variables. Check the validity of the data (especially the *temp* variable) and repeat the previous script stages if necessary. |

**STEP 11FA: Count objects in the '*ring areas'***

1. Run the 'FOCAL ADHESIONS ADD-ON | COUNT NUMBER OF OBJECTS CUTTING THROUGH BANDS OF GIVEN DIAMETER' section to analyze the distribution of objects within the previously defined '*ring areas*' – concentric bands estimated consecutively from the cell border (defined by the *cell ROI*) toward the cell interior.

**NOTE:** Larger objects intersecting multiple '*ring areas*' will be counted in the total for each ring they pass through.

**TROUBLESHOOTING:** The possible errors are the same as those that may occur during the initial analysis of emission intensity within the selected '*ring areas*'. Refer to the descriptions of errors E15-E18 and warnings W9 and W10 in the *TROUBLESHOOTING* subsection of *STEP 6FA: Analysis of emission intensity within concentric bands of a specified diameter ('ring areas')* in the manual.

**STEP 12FA: Transfer from pixel to real size units**

1. Run the 'FOCAL ADHESIONS ADD-ON | CHANGE PX DISTANCES TO REAL SIZE UNITS' section to convert pixel-based measurements used throughout the script into real-world units, either retrieved from image metadata or manually provided by the user via the console window.

**NOTE:** If this stage is omitted, the script will proceed normally; however, all measurements will remain in pixel units and will be saved as such in the final spreadsheet.

**TROUBLESHOOTING**

| Number of error/warning | Error / warning prompt | Error / warning description | Recommended solution |
|---|---|---|---|
| E22 | FOCAL ADHESIONS ADD-ON \| ERROR: Transfer from distances in pixels to real size failed. Check data and try again. | The conversion to real-world units was unsuccessful. In case of failure, the data will remain in pixel units. | Carefully verify the data integrity following a failed unit conversion (with particular attention to the *temp* variable). If the entire dataset remains in pixel units, ensure that the *px_size* variable contains valid numeric data – either manually provided by the user or extracted from the image metadata. If not, input the correct value via the console window and rerun the section. The user should also validate the correctness of all other input variables used in the *talPxTransfer* method. If the data within the *temp* variable has been only partially converted, this may indicate structural |

| | | | |
|---|---|---|---|
| | | | changes in the object property storage caused by previous script operations. In such cases, the only reliable solution is to rerun the entire analysis. Users are encouraged to contact the authors for data recovery support. Copies of the workspace variables, original image files, and any result spreadsheets (if generated) will be required as source materials. |
| W16 | FOCAL ADHESIONS ADD-ON \| WARNING: This operation cannot be called if you have already performed one transfer to real size units. | This warning appears if the unit conversion has already been performed, preventing duplicate conversion. | No action is required. This warning indicates that a user action was blocked to prevent unintended script behavior. The previously converted units will remain unchanged. |

**STEP 13FA: Results display and saving**

1. Run the 'FOCAL ADHESIONS ADD-ON | RESULTS DISPLAY' section to display and save all generated maps, overlays, and charts. The adjusted images, visual maps, and charts will be saved in the source folder of the input image used for the focal adhesions add-on, in both TIFF and FIG formats.

**NOTE:** The user is advised to store the input image in a separate directory to facilitate data management after file saving.

**TROUBLESHOOTING**

| Number of error/warning | Error / warning prompt | Error / warning description | Recommended solution |
|---|---|---|---|
| E23 | FOCAL ADHESIONS ADD-ON \| ERROR: Results cannot be fully displayed. Check data. | The results cannot be fully displayed and saved. While the display failure will not affect the data itself, it will result in only a partial saving of the results. | Identify which image caused the issue with data display and saving. The problematic image may not be displayed or saved, or a blank window may appear in its place after the last successfully displayed image. Then, verify the validity of the data associated with the image that failed to display. If the data is correct, check whether the source directory is being used by another process, which could be preventing the saving operation and blocking the script from continuing after the first image is displayed. Ensure that all applications using the directory are closed, and close the directory window itself before trying again. If the failure occurs again, manually save the results using the display windows. |

**STEP 14FA: Saving results to spreadsheet**

1. Run 'FOCAL ADHESIONS ADD-ON | SAVE RESULTS TO SPREADSHEET' to export data to the spreadsheet (.xlxs format), which will contain:
   a. Full object parameter data, including: area, centroid, circularity, perimeter, major axis length, minor axis length, maximum/minimum dimension ratio, orientation, intensity centroid, mean intensity, minimum intensity, maximum intensity, an indicator of whether the object was manually added by the user, and whether it is located above the cell nucleus area.
   b. Data calculated for the '*ring areas*', including: intensity values within each concentric band and the number of objects intersecting each band of the specified width.
   c. Workspace data, including final variable values and result images in MAT-format file.

**NOTE:** You can run this section multiple times, including after label or ROI corrections. In such cases, the data will be overwritten and previous results will be replaced.

**NOTE 2:** If the analysis must be interrupted due to external circumstances, the user can store the workspace variables and resume the analysis later by using the following command: save(strcat(filepath, 'BinaryMaps-', save_default_name));. The temporary workspace will be saved in the directory containing the source image. To resume the analysis, the user can reload the workspace using MATLAB's *load* function by specifying the correct file path (refer to the MATLAB documentation for syntax and proper file path usage). Once loaded, the user can continue the analysis from the point at which it was previously saved.

**Performance considerations**

1. Selecting the DnCNN-based denoising method (*denoising_flag* = 1) may significantly increase processing time. If processing time or computational resources are a concern, choosing an alternative method or disabling denoising may still yield satisfactory results.
2. Processing time increases with image size and complexity.
3. If performance slows down during multiple ROI modifications within a single session or after merging multiple labels in one step, the user is encouraged to accept the current changes and continue with further additions, deletions, or mergings in subsequent steps. It is recommended to process a smaller number of objects across multiple sessions rather than applying all changes within a single editing session.

**Recommended Parameter Adjustments for Common Scenarios**

1. For dim or low contrast images:
   a. Ensure *contrast_flag* = 2 for manual control and decrease *lh_lim_tal* first value to ~ 0.05 and second value to ~ 0.2 - 0.5
   b. Set *binarization_met* = 1 and decrease *segm_threshold* to ~ 0.35
2. For noisy images:
   a. Enable *denoise_flag* = 1 or *denioise_flag* = 2 (for faster performance)
   b. Turn on background subtraction (*rb_flag* = 1) and use *rb_method* = 1; for faster performance consider testing the small structural element size at first (*se*)
   c. Set *filt_flag* = 2 (bilateral filtering)
   d. Consider increasing minimum area threshold slightly
3. For dense adhesion structures:
   a. If background subtraction is enabled choose rolling ball radius method (*rb_flag* = 0) and consider increasing the *rb_thresh* value to prevent removal of relevant structures; if processing performance is too slow, try slightly decreasing the *rb_thresh* value
   b. May need to increase *ratio_lvl* if structures appear interconnected
4. For analysis of smaller, round objects (such as nascent adhesions):
   a. Decrease *obj_area_thresh_itsty, obj_area_thresh_circ_itsty* and *obj_area_thresh* to include smaller objects. Remember that this may lead to inclusion of unwanted noise or artifacts in final classification
   b. Try running the analysis with denoising disabled (*denoise_flag* = 0); small artifacts should still be excluded by the classifier and filtering steps. If you choose to enable denoising using the morphological-based approach (*denoise_flag* = 2), use a small structural element size (*se*)
   c. Consider setting *binarization_met* = 2 (adaptive segmentation with high sensitivity) to better detect dim, small structures
   d. Decrease *itsty_lvl* to ~0.5-0.6 to include dimmer nascent adhesions
   e. Lower *circ_param* slightly (below 0.7) to accept more circular objects

Tab. S1. The list of parameters and flags used to enable or disable specific operations, available for user adjustment.

| VARIABLES | DEFAULT VALUE | RANGE | DESCRIPTION |
|---|---|---|---|
| **Essential Control Flags** | | | |
| FA | 1 | {0,1} | Controls the adhesion structure analysis module (*focal adhesions* add-on). When set to 1, focal adhesion segmentation and feature extraction are enabled. Set to 0 to disable the module. This add-on requires a single image channel to function properly. |
| colocalization | 1 | {0,1} | Controls the colocalization analysis module (*colocalization* add-on). When set to 1, it enables the generation of colocalization maps and the calculation of correlation coefficients between two image channels. Set to 0 to disable the module. This add-on requires two image channels to function properly. |
| **Method execution flags** | | | |
| auto_roi_flag | 1 | {0,1} | Specifies the *cell ROI* selection method: 0 – manual ROI selection, where the user draws the cell outline; 1 – automatic ROI detection. Automatic detection performs best with well-defined cell boundaries and high contrast. It is recommended to begin with automated *cell ROI* detection. |
| contrast_flag | 0 | {0,1,2} | Controls image contrast settings (applied independently of visualization for *cell ROI* / *segmentation ROI* selections): 0 – applies automatic contrast adjustment based on the image histogram; 1 – applies automated Otsu-based contrast limits for image analysis; 2 – applies user-defined intensity limits specified in *lh_lim_tal* or *lh_lim_coloc* (depending on the add-on choice) for analysis. |
| contrast_visual_flag | 0 | {0,1} | This option allows the user to apply custom contrast limits (user-defined only) for visualization during *cell ROI* or *segmentation ROI* selection, separate from those used for calculations. This is useful, for example, when enhanced contrast is needed for manual selection, but should not influence analytical computations: 0 – contrast settings defined by automated adjustment or specified in *lh_lim_tal* or *lh_lim_coloc* will be used for both calculations and visualization; 1 – contrast settings defined in *lh_lim_tal_user* or *lh_lim_coloc_user* will be used for visualization only, while those in *lh_lim_tal* or *lh_lim_coloc* will be used for calculations. |
| denoise_flag | 1 | {0,1,2} | Controls the selection method for denoising algorithms: 0 – off; 1 – DnCNN-assisted |

| | | | |
|---|---|---|---|
| | | | denoising (may be computationally intensive and time-consuming on older devices); 2 – morphological operations-based denoising. |
| rb_flag | 1 | {0,1} | Controls background subtraction: 0 – no background subtraction; 1 – applies background subtraction using the method specified by the value of the *rb_method* variable. |
| filt_flag | 2 | {0,1,2,3} | Selects the filtering method: 0 – bilateral filtering with high smoothing; 1 – Gaussian filtering; 2 – bilateral filtering with moderate smoothing; 3 – Savitzky–Golay filtering. Bilateral filtering (option 2) is recommended as the default option for initial use, as it effectively reduces noise while preserving edge details. |
| manualcorr_flag | 0 | {0,1} | Allows the user to specify whether manually added objects should be included in the label map if they do not meet the classification criteria: 0 – all user-added objects override the classifier conditions and are included in the final label map; 1 – user-added objects undergo classification and are excluded from the label map if they do not meet the classification criteria. |
| ring_hole_flag | 0 | {0,1} | Allows for the choice if holes in cell body (if exist) should be enveloped by the '*ring areas*', or ignored and treated as the cell body during the extraction of '*ring areas*': 0 – holes in the cell body will be ignored and included in the 'ring areas', 1 – holes in the cell body will be enveloped by the '*ring areas*'. |
| **Method variant selection** | | | |
| binarization_met | 3 | {0,1,2,3} | Selects segmentation method: 0 – adaptive segmentation (low sensitivity), 1 – global thresholding with user-defined threshold (*segm_threshold*), 2 – adaptive segmentation (high sensitivity), 3 – global segmentation with threshold defined via Otsu's method. |
| rb_method | 0 | {0,1} | Selects the background subtraction method when rb_flag = 1: 0 – rolling ball radius method; 1 – morphological operations-based method. |
| **Image processing parameters** | | | |
| se | Shape: sphere<br>Size: 2 | Min: 1<br>Max: image size (depends on image smaller dimension)<br>Recommended max: 50 | Sets the size of the structuring element used for morphological operations by modifying the 'se = strel('sphere', 2)' clause, where the first input parameter specifies the shape of the structuring element and the second defines its size (diameter). Refer to the MATLAB documentation for the *strel* function for detailed syntax. Smaller values preserve fine details, while larger values emphasize broader structures. It is recommended to use the |

| | | | |
|---|---|---|---|
| | | | spherical shape and adjust only the size. Note that larger structuring elements may significantly increase processing time. Excessively large values may cause the software to malfunction or throw exceptions. It is advised to begin with small values and incrementally increase the size, monitoring the results at each step. For practical and computational reasons, it is recommended not to exceed a radius of 100 pixels. |
| gauss_sd | 2 | Min: 0 (delta function)<br>Recommended min: 0.5<br>Max: image size (depends on the image's smaller dimension)<br>Recommended max: 15 | Controls the standard deviation (σ) of the Gaussian filter when *filt_flag* = 1. Higher values result in stronger smoothing. While there is no strict maximum limit, it is recommended not to exceed *gauss_sd* = 20 for practical performance reasons, and not to use values below *gauss_sd* = 0.3 to avoid unintended script behavior. |
| rb_thresh | 2 | Min: 1<br>Max: image size (depends on the image's smaller dimension)<br>Recommended max: 50 | Sets the radius (in pixels) for rolling ball background subtraction when *rb_method* = 0. It is recommended to choose a radius comparable to or larger than the diameter of the objects of interest to prevent them from being removed as background. For practical and computational reasons, it is recommended not to exceed a radius of 100 pixels. |
| **Segmentation & classification parameters** | | | |
| segm_threshold | 0.5 | <0,1> | Sets the segmentation threshold for the global segmentation method when an automatically chosen threshold is not used (*binarization_met* = 1). |
| ***Focal adhesions* add-on parameters** | | | |
| ch_tal | Green channel: 2 | {1,2,3} | Selects the channel for adhesion structures analysis in RGB images: 1 – red, 2 – green, 3 – blue. |
| lh_lim_tal | [0.1, 0.9] | <0;1> for each vector coordinate | Sets the contrast limits for the chosen image channel. The two-element vector defines the lower and upper intensity scaling limits: [lower limit, upper limit]. Increasing the lower limit results in more pixels being assigned to the image background, while decreasing the upper limit enhances the visibility of darker objects by brightening them to the level of maximal allowed image intensity. |
| lh_lim_tal_user | [0.1, 0.9] | <0;1> for each vector coordinate | Sets the contrast limits for the selected image channel. The two-element vector [lower limit, upper limit] defines the lower and upper intensity scaling limits, similarly to *lh_lim_tal*. These limits are applied only to the image used for visualization during manual ROI |

| | | | |
|---|---|---|---|
| | | | adjustments by the user (when *contrast_visual_flag* = 1). |
| ring_diameter_µm | 1 | Min: > 0<br>Max: cell diameter (depends on cell size)<br>Recommended max: 2 | Sets the width of the '*ring areas*' – consecutively drawn concentric bands starting from the cell border and extending toward the cell interior or across internal holes (depending on the value of the *ring_hole_flag* variable) – expressed in micrometers (µm). The maximum ring width should preferably be less than half of the shortest cell 'diameter'. Larger values may result in only a single ring encompassing the entire cell area. The appropriate ring width should be determined by the user based on the size of the analyzed cell. |
| obj_area_thresh_itsty | 100 | Min: 1<br>Max: image size (depends on image smaller dimension)<br>Recommended max: depends on the type of cells / to be decided by user | Specifies the minimum area (in pixels) required to accept objects with intensity above the *itsty_lvl* threshold (range: <0,1>). Filters out small bright specks that are likely noise rather than actual adhesion structures. |
| obj_area_thresh_circ_itsty | 110 | Min: 1<br>Max: image size (depends on image smaller dimension)<br>Recommended max: higher than *obj_area_thresh_itsty*, depends on the type of cells / to be decided by user | Specifies the minimum area (in pixels) required to accept circular objects with intensity above the *itsty_lvl* emission intensity threshold (range: <0,1>). This separate threshold is used to more effectively exclude circular objects, which are less likely to be classified as focal adhesions. |
| obj_area_thresh | 150 | Min: 1<br>Max: image size (depends on image smaller dimension)<br>Recommended max: higher than *obj_area_thresh_itsty*, depends on the type of cells / to be decided by user | Specifies the minimum area (in pixels) for accepting any object, regardless of its emission intensity or shape. This value should be set based on the expected size of larger adhesion structures that may appear with low intensity due to specific staining conditions. |
| circ_param | 0.7 | <0,1> | Specifies the circularity threshold for object classification. Lower values allow more circular objects to be accepted as adhesion structures in the final *segmentation ROI* after classification, while higher values result in greater exclusion of circular objects from the final map. A starting value of *circ_param* = 0.7 is recommended, as set by default. |
| itsty_lvl | 0.65 | <0,1> | Specifies the intensity threshold for excluding objects based on intensity criteria, in conjunction with object area (as defined by the *obj_area_thresh_itsty* and |

| | | | |
|---|---|---|---|
| | | | *obj_area_thresh_circ_itsty* variables). Objects with mean intensity below this threshold may be excluded depending on their area (only the largest objects with intensity below the *itsty_lvl* threshold will be retained in the final *segmentation ROI* after classification, as determined by the *obj_area_thresh* variable value). |
| itsty_lvl_ROI | 0.35 | <0,1> | Specifies the minimum intensity threshold for pixel inclusion as part of the neighborhood during automatic *cell ROI* detection. Lower values result in the inclusion of more low-intensity regions within the *cell ROI* (recommended if parts of the cell are being omitted). Higher values lead to greater exclusion of low-intensity pixels.<br>NOTE: This variable does not operate as a strict emission intensity threshold like *itsty_lvl*, as pixel neighborhood is also taken into account. Users are encouraged to test different values and observe the resulting behavior to achieve the desired outcome if the default setting does not yield satisfactory results. |
| ratio_lvl | 2 | Min: 1 (would be related with circular objects or single pixels)<br>Recommended min: 1.5<br>Max: image size (depends on image smaller dimension)<br>Recommended max: 3.0 | Specifies the object dimensional ratio threshold for elimination based on shape. Objects are excluded if the ratio of their largest to smallest dimension is below *ratio_lvl*, in combination with minimum area and emission intensity criteria. This threshold helps distinguish elongated (fibrillar) structures from more circular ones. It is recommended to start with lower values (e.g., *ratio_lvl* = 2.0) and increase incrementally, to avoid excluding less elongated objects that may still qualify as focal adhesion structures. |
| ***Colocalization* add-on parameters** | | | |
| ch_coloc | Red & green channels: [1, 2] | {1,2,3} | Selects the channels for colocalization analysis in RGB images, where: 1 – red, 2 – green, and 3 – blue. The selection is defined as a two-element vector specifying the image channels to be analyzed by the *Colocalization* add-on: [first_protein_channel, second_protein_channel]. These values correspond to the channels selected by the user during image input. For example, if the vector is set to *ch_coloc* = [3, 1], the first selected channel will be displayed in the blue layer and the second in the red layer of the resulting RGB image. This allows the user to customize the visualization order regardless of the original staining colors. |
| lh_lim_coloc | [0.1, 0.9, 0.1, 0.9] | <0;1> for each vector coordinate | Sets the contrast limits for colocalization analysis, similar to the *lh_lim_tal* parameter used in the *focal adhesions* add-on. The contrast is defined using a four-element vector: [lower limit for first channel, upper limit for first channel, lower limit for second channel, |

| | | | |
|---|---|---|---|
| | | | upper limit for second channel]. These values are used for independent intensity scaling of the two selected image channels. Increasing the lower limit results in more pixels being assigned to the image background, while decreasing the upper limit enhances the visibility of darker objects by brightening them toward the maximum allowable image intensity.<br>NOTE: In the case of automatic contrast adjustment, the two channels are scaled independently. There is no option to apply identical automatic settings to both channels; this is only possible when contrast limits are set manually (*contrast_flag* = 2). |
| lh_lim_coloc_user | [0.1, 0.9, 0.1, 0.9] | <0;1> for each vector coordinate | Sets the contrast limits for the selected image channels, similar to the *lh_lim_col* parameter. The four-element vector [lower limit for first channel, upper limit for first channel, lower limit for second channel, upper limit for second channel] defines the lower and upper intensity scaling limits for both channels. These limits are applied only to the image used for visualization during manual ROI adjustments (when *contrast_visual_flag* = 1). |
| labels_cbar | For default green and red channels: {'G intensity MAX', 'Equal intensity', 'R intensity MAX'} | Any text consisting of three separate phrases enclosed in single quotation marks (' ') and separated by commas | Labels for the colorbar in colocalization maps. Customize these to match your protein names for clearer visualization. |
